# Covalent bond formation caught in a LOV photoreceptor

**DOI:** 10.64898/2026.08.02.742302

**Authors:** Bence Olasz, Guillaume Gotthard, Probal Nag, María González-Viegas, Sandra Mous, Anna Koczurowska, Philip J. M. Johnson, Tokushi Sato, Madan Kumar Shankar, Maximilian Wranik, Aleksandra Solecka, Diogo Melo, Ran Ke, Daniel James, Karol Nass, Pit Langner, Joana Valerio, Antonia Furrer, Dardan Gashi, Raphael de Wijn, Tomas Popelar, Magdalena Pachota, Magdalena Wisniewska, Thomas Dietze, Juncheng E, Ayesha Asghar, Dmitrii Zabelskii, Fabian Trost, Faisal H. M. Koua, Peter Smyth, Egor Sobolev, Chan Kim, Dmitry Ozerov, Romain Letrun, Oleksii Turkot, Katerina Doerner, Johan Bielecki, Huijong Han, Florian Dworkowski, Claudio Cirelli, Gebhard F. X. Schertler, Joachim Schulz, Camila Bacellar, Joerg Standfuss, Richard Bean, Chris Milne, Joachim Heberle, Igor Schapiro, Przemyslaw Nogly

## Abstract

Light-oxygen-voltage (LOV) domains are blue-light photoreceptors of plants, algae and fungi, and among the most widely used tools in optogenetics. They switch on by forming a covalent thioether bond between a conserved cysteine and their flavin chromophore, in a reaction that needs a proton to cross from the cysteine to the flavin through a pocket containing essentially no water. Its mechanism has been debated for two decades^1^, and because the chemistry is over within a microsecond its elementary steps have stayed hidden. Here we combine 10 time-resolved serial femtosecond crystallography snapshots and infrared spectroscopy with QM/MM calculations to resolve the entire sequence of events at 1.4 Å resolution: from excited-state distortion of the flavin ring (10–100 ps), through hydration of a surface channel (10 ns) and a single ordered water reaching the active site as the reactive cysteine shifts between its conformations (100–500 ns), to the thioether bond itself, caught half-formed at 1 µs (half the molecules reacted, half still poised) and complete at 10–100 µs. That water bridges the cysteine and the flavin and shuttles the proton, lowering the barrier from ∼35 to ∼15 kcal/mol and accelerating the reaction by roughly fourteen orders of magnitude (without it, the half-life would be ∼237,000 years), then departs before the bond forms. Proteins can therefore hydrate a dehydrated active site transiently and on demand to overcome otherwise prohibitive reaction barriers, a catalytic strategy that reaches well beyond photoreceptors.

## Introduction

Phototropins are blue-light photoreceptors found in green algae and all major land plant lineages^2,3^, where they control phototropism^4^, stomatal opening^5^, and chloroplast movements^6^. In *Chlamydomonas reinhardtii (Cr)*, phototropin additionally regulates the sexual life cycle^7^, gene expression^8^, photosynthesis^9^, and eyespot development^10^. These receptors contain tandem N-terminal light-oxygen-voltage (LOV) domains that bind a flavin mononucleotide (FMN) chromophore and undergo a photocycle upon blue-light absorption, forming a covalent thioether adduct between a conserved cysteine and the FMN C4a^11^. The founding LOV crystal structure placed this cysteine 4.2 Å from C4a and first proposed the adduct model^12^.

The photocycle of the *Cr*LOV1 domain has been characterized spectroscopically: photoexcitation produces a singlet excited state that converts via intersystem crossing (τ = 3.5 ns) into two triplet subspecies, LOV_715a_ and LOV_715b_, which decay into the thiol adduct (LOV_390_) with time constants of 800 ns and 4 µs, respectively^13,14^(Fig. 1). The adduct spontaneously reverts to the dark state in ∼200 s^13,14^ot (Supplementary Fig. 1).

**Fig. 1:**
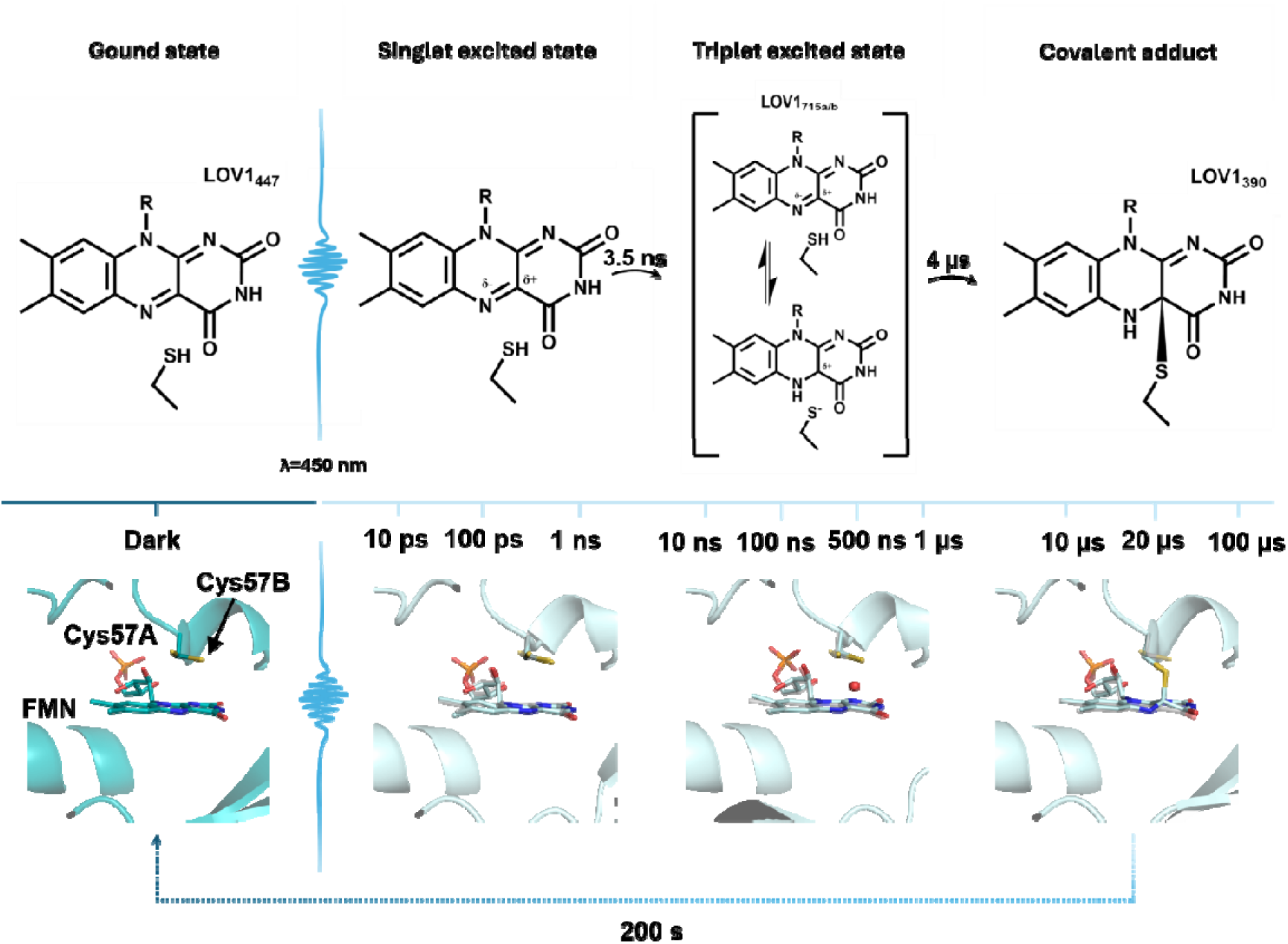
TR-SFX resolves the *Chlamydomonas reinhardtii* LOV1 photocycle from femtosecond light absorption to microsecond covalent adduct formation, identifying three mechanistic stages. **Upper**. The *Cr*LOV1 photocycle. Blue-light absorption (λ = 447 nm) promotes the FMN chromophore from the ground state (LOV1_447_) to a singlet excited state (lifetime 3.5 ns), which undergoes intersystem crossing to two triplet subspecies (LOV1_715a_, LOV1_715b_). These decay to the covalent thioether adduct (LOV1_390_) with time constants of 800 ns and 4 µs, respectively^13,14^; the adduct reverts thermally to the ground state in ∼200 s. Chemical structures show the FMN isoalloxazine ring and reactive Cys57 at each spectroscopic state. **Middle.** Timeline of all TR-SFX time delays captured in this study (10 ps to 100 µs), positioned within the photocycle kinetics to indicate which intermediates are structurally resolved. **Lower.** Representative structural snapshots of the FMN binding pocket at 100 ps, 500 ns, and 20 µs, each superimposed on the dark-state reference structure (grey; PDB: 8QI8; 1.4 Å). The three panels illustrate the three principal mechanistic stages: isoalloxazine ring distortion (100 ps), transient active-site hydration (500 ns), and fully formed covalent adduct (20 µs). Light-state structures are shown in light blue.

Static crystal structures of *Cr*LOV1 have captured the dark and late light states^15^, and time-resolved serial synchrotron crystallography (TR-SSX) resolved millisecond rearrangements, including C-terminal shifts linked to signal transduction^16^. However, because covalent adduct formation completes within ∼1 µs, these millisecond snapshots captured the protein only after the critical photochemistry had already occurred.

The elementary steps coupling light absorption to covalent bond formation, including the debated nature of the proton-transfer event preceding thioether bond formation at C4a^1^, have therefore remained structurally unresolved. Solvent isotope exchange has established proton transfer as an essential, rate-limiting step of adduct formation: the reaction is ∼5-fold slower in DOO than in HOO, an effect abolished when the reactive cysteine is removed^17^. Yet how this kinetically forbidden proton transfer proceeds across a largely dehydrated chromophore pocket has remained an open question.

Here, we combine TR-SFX with hybrid QM/MM calculations and time-resolved infrared spectroscopy to resolve the *Cr*LOV1 photocycle intermediates from 10 ps to 100 µs, including covalent adduct formation. Transient hydration of the chromophore environment recruits a single ordered water molecule between Cys57 and FMN N5, enabling the kinetically forbidden proton transfer that precedes covalent bond formation. This transient catalytic hydration — accelerating the reaction by ∼14 orders of magnitude — reveals a fundamental strategy by which enzymes may overcome energy barriers to proton shuttling in dehydrated active sites.

## Results

### FMN distortions characterize the early photocycle

*Cr*LOV1 crystals diffracting to 1.4 Å resolution enabled capture of ultrafast photocycle dynamics across time delays from 10 ps to 100 µs by TR-SFX (Fig. 1, Extended Data Table 1). Photoexcitation triggers immediate out-of-plane distortions of the FMN isoalloxazine ring. At 10–100 ps, positive difference electron density appears near FMN N5 (+3.8 σ; Fig. 2a), yielding a 0.3 Å out-of-plane displacement. Excited-state polarization of the isoalloxazine ring — with increased electron density at N5 (δ−) and electrophilicity at C4a (δ+) — is established in related LOV domains^18^ which accounts for the observed displacement by increasing single-bond character at N5 and disrupting sp^2^ planarity.

**Fig. 2:**
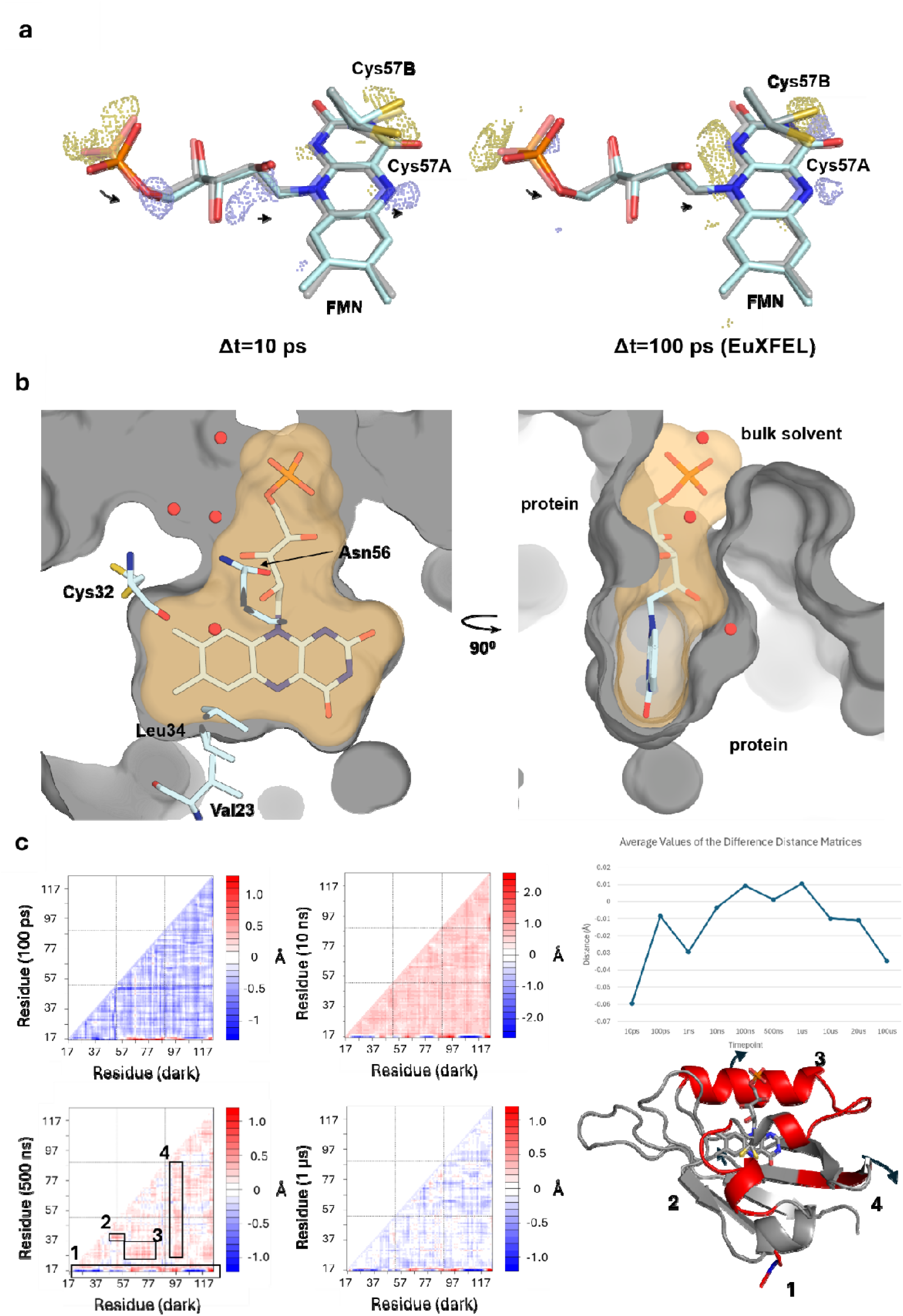
Picosecond phosphoribityl tail contraction and global Cα analysis reveal a protein-wide contraction–expansion cycle that precedes active-site hydration. (**A**) Fo–Fo difference Fourier maps at Δt = 10 ps (left) and 100 ps (right; collected at the European XFEL), contoured at ±3.0 σ (positive, blue; negative, gold). At 10 ps, density marks a 0.5 Å outward phosphate displacement (+4.6/−5.4 σ) and a 0.3 Å C1′ shift (+3.3 σ), persisting at 100 ps (+3.6/−4.6 σ). The 100 ps result is reproduced across XFEL facilities and delivery methods (Extended Data Fig. 1). Light state, light blue; dark reference, grey; 1.4 Å; Cys57 conformations A/B. (**B**) Surface of the *Cr*LOV1 FMN pocket (water, red spheres); the phosphoribityl tail is exposed to bulk solvent, allowing transient water ingress. (**C**) Cα distance matrices at 100 ps, 10 ns, 500 ns and 1 µs versus the dark state (blue, contraction; red, expansion), with mean Cα displacement as a line graph. Contraction in the singlet state (100 ps) switches to expansion at triplet formation (10 ns), then relaxes by 500 ns–1 µs. Inset (500 ns), coloured by per-residue displacement, marks the FMN-flanking, phosphate-tail, water-gate and Gln120 regions as the zones of greatest shift before adduct formation.

Concomitant with ring deformation, the FMN phosphoribityl tail — which projects into bulk solvent (Fig. 2b) — contracts toward the FMN ring (Fig. 2a), defining a potential structural conduit for transient water ingress into the chromophore pocket. At 10 ps, the phosphate group displaces by 0.5 Å (+4.6/−5.4 σ) and C1□ shifts by 0.3 Å (+3.3 σ; Fig. 2a). At 100 ps, these changes persist with reduced amplitude (+3.6/−4.6 σ). Independent experiments at two XFEL facilities with different delivery methods reproduced the 100 ps structural changes (Fig. 2a, Extended Data Fig. 1b, 1d).

Ring planarity largely recovers by Δ*t* = 10 ns, coincident with the singlet-to-triplet transition^13,14^ (Supplementary Fig. 1, Supplementary Fig. 2). Residual out-of-plane shifts persist into the nanosecond regime: an elevation at C8/C8M gains difference density support at 100 ns (+3.5 σ; Supplementary Fig. 3b), and a shift near C2/O2 — close to the reactive center of the ring — emerges at 500 ns (+4.1 σ; Supplementary Fig. 3c, Supplementary Fig. 4a).

Beyond these residual ring distortions, global backbone analysis resolves coordinated whole-protein motions across the photocycle. Cα distance analysis across all TR-SFX time delays (Extended Data Fig. 2) reveals a global protein contraction in the singlet state (100 ps) that switches to expansion coincident with triplet formation (10 ns) and then relaxes toward the dark-state configuration by 500 ns (Fig. 2c). At 500 ns, the residues flanking FMN, the phosphate tail, and the C-terminal region near Gln120 show the greatest persistent displacement, identifying the structural zones most affected immediately before adduct formation (Fig. 2c).

### Active-site hydration precedes adduct formation

The ultrafast phosphoribityl contraction dissipates in the early nanoseconds (Extended Data Fig. 1c), though continued tail displacement persists at 100 ns (−3.9 σ; Extended Data Fig. 3b). By 500 ns, the phosphate group shifts 0.7 Å toward the sidechain of Arg58 (+3.9/−6.9 σ; Extended Data Fig. 3c, Supplementary Fig. 5b). Arg58 and Arg74, flanking the FMN phosphate at the solvent interface, show increased mobility with displacements up to 2.4 Å (Supplementary Fig. 6). Main-chain shifts of 0.24–0.34 Å extend from Asn56 to Gln61 (up to +4.1/−6.1 σ at 500 ns; Fig. 2c, Extended Data Fig. 4), with concomitant sidechain movements of Asn56 and Cys57. Asn56, which hydrogen-bonds the phosphate tail in the dark state, shifts by 2.8 Å (+3.5 σ in F_500ns_–F_dark_), underscoring the extent of interfacial disorder preceding adduct formation. A light-induced increase in disorder at the phosphate tail across time points is summarized in Extended Data Fig. 5.

The key event on the nanosecond timescale is transient hydration of the chromophore pocket. The dark-state structure of *Cr*LOV1 is largely dehydrated, with only a single distant water molecule (Wat1, 3.8 Å from FMN) near the Cys57 main chain (Fig. 3a). After photoactivation, additional solvent molecules penetrate the protein core from Δ*t* = 10 ns (Fig. 3a, Supplementary Fig. 7). This influx coincides with the global backbone expansion resolved at 10 ns (Fig. 2c), indicating that transient loosening of the protein accompanies solvent entry into the core. This hydration culminates in the appearance of a new water binding site, Wat134, at 100 ns between the reactive thiol of Cys57 (2.7 Å) and the N5 atom (2.4 Å; +3.2 σ in F^100ns^–F^dark^) with partial occupancy at this stage (Fig. 3a). Wat134 persists at 500 ns with increased occupancy (+3.5 σ; Fig. 3a), refining into well-defined density with hydrogen-bonding distances to FMN N5 and Cys57 (Fig. 3c). Weak positive density (+3.1 σ) is already detectable in F^10ns^–F^dark^ at the site later occupied by Wat134, but its proximity to a neighboring residue precludes reliable refinement. The appearance of Wat134 immediately before bond formation implicates transient active-site hydration as the pathway for proton transfer from Cys57 to FMN N5.

**Fig. 3:**
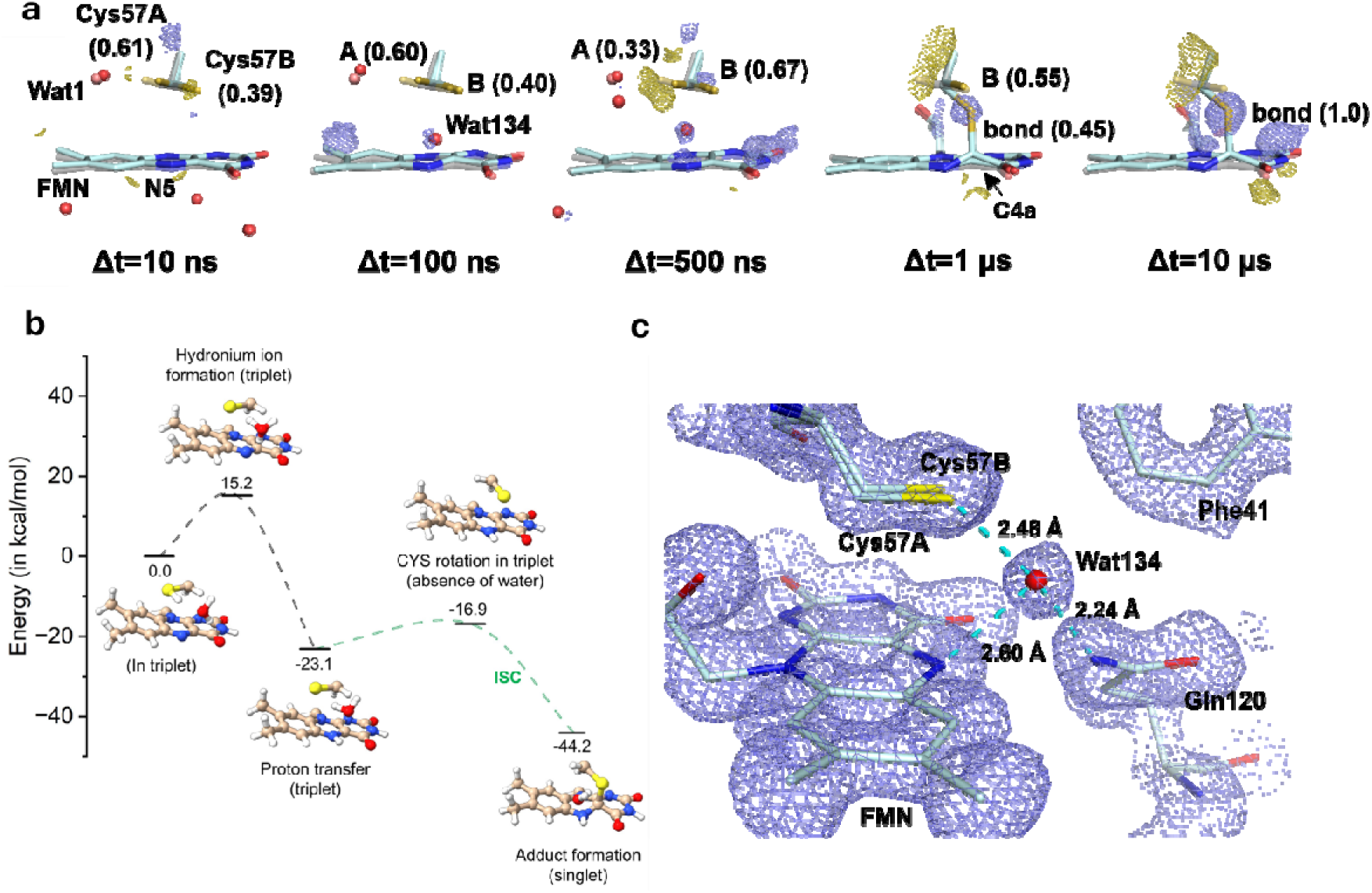
A transiently ordered water molecule (Wat134) appears between Cys57 and FMN N5 enabling proton transfer and thioether bond formation by 10 µs. (**A**) Fo–Fo difference maps (±3.0 σ; positive, blue; negative, gold; 1.4 Å) at five delays from triplet to adduct. At 10 ns the site resembles the dark state, weak density (+3.1 σ) marking the future Wat134 site. At 100 ns Wat134 appears at partial occupancy (+3.2 σ), 2.7 Å from the Cys57 thiol and 2.4 Å from N5. At 500 ns occupancy rises (+3.5 σ), Cys57 redistributes from 0.52:0.48 in resting state to 0.33:0.67 (A:B), and C4a sp^2^→sp^3^ rehybridisation begins. At 1 µs conformation A is gone, a partial bond forms, and Wat134 is absent. At 10 µs the thioether bond is complete (+7.6/−7.9 σ; 109° angles). Light state, light blue; dark, grey; Wat134, red sphere; Cys57 A/B. (**B**) QM/MM profile for the favoured pathway (P-III): water-mediated transfer via a hydronium-ion transition state, 15.2 kcal/mol, versus 35.3 kcal/mol without Wat134; cysteine rotation then needs ∼6 kcal/mol (full diagram, Supplementary Fig. 9). (**C**) Active site at 500 ns: Wat134 hydrogen-bonds N5 and nearby residues; 2Fo–Fc map (blue, 1.5σ).

### A solvent access pathway analogous to the *As*LOV2 water gate

The arrival of Wat134 in a largely dehydrated core raises the question of which pathway channels bulk solvent to the buried FMN pocket. Hydrophobic residues Val103, Pro105, and Phe116 obstruct the potential path along the phosphoribityl tail in *Cr*LOV1. Nevertheless, mutant studies show that the core is accessible to solutes, as demonstrated by methylmercaptan adduct formation^19,20^.

In the dark state, *Cr*LOV1 presents a closed surface channel (Fig. 4) lined by Thr21 and Gln120, with only a peripheral water near its mouth; this tunnel is structurally equivalent to the *As*LOV2 water gate (Asn414/Gln513) proposed based on mutagenesis and dark-state recovery kinetics^21^. Despite the Asn→Thr substitution, our time-resolved structures show that this tunnel becomes transiently solvent-accessible upon photoactivation. At Δ*t* = 10 ns, a transient water molecule (Wat49) fills the channel (Fig. 4c), plausibly delivering catalytic Wat134 to the active site. When Wat134 disappears at 1 µs upon covalent adduct formation, Wat49 reappears at the water gate site (Fig. 4f), defining a return route for active-site dehydration. These observations provide the first time-resolved structural evidence for a LOV domain water gate and reveal that *Cr*LOV1 operates a dynamic hydration–dehydration cycle coupling solvent access to covalent bond formation and its stabilization (Supplementary Fig. 8). Together, these dynamics position a water molecule at the reactive centre precisely when adduct chemistry occurs, motivating a quantitative assessment of its catalytic contribution.

**Fig. 4:**
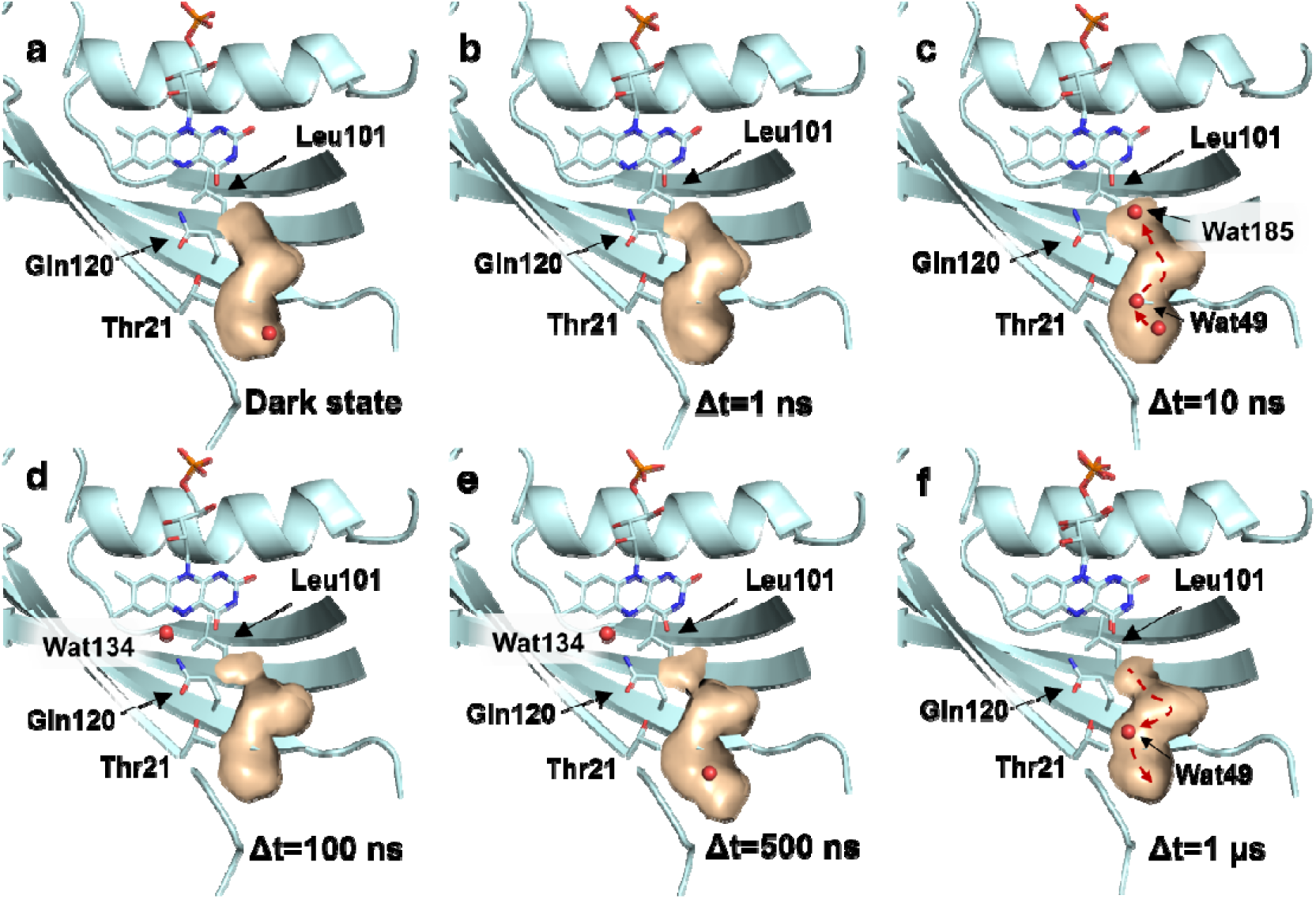
Time-resolved crystallography reveals the *Cr*LOV1 water gate in operation: Wat49 enters the chromophore pocket at 10 ns and retreats at 1 µs, coupling active-site hydration to photocycle progression. (**A**) Dark-state tunnel interior (orange surface) with gate residues Thr21 and Gln120 shown as labelled sticks. Thr21 and Gln120 are the *Cr*LOV1 structural equivalents of residues identified as water gate in *As*LOV2 Asn414 and Gln513, respectively. In the dark state, the channel is closed; only a surface water molecule is present near the gate. (**B**) At Δ*t* = 1ns the tunnel interior lacks water molecules entirely. (**C**) At Δ*t* = 10 ns, water molecules Wat49 and Wat185 occupies the tunnel interior, indicating solvent ingress coincident with triplet-state formation and the onset of active-site hydration. (**D**) At Δ*t* = 100 ns the interior water molecules from previous probed timepoint have disappeared. (**E**) At Δ*t* = 500 ns the water molecule near the surface is observed. (**F**) At Δ*t* = 1 µs, following covalent bond formation and the catalytic departure of Wat134, Wat49 reappears at the water gate, indicating a return pathway for active-site dehydration. Panels **C** and **F** present the first direct time-resolved structural evidence for a dynamic nature of LOV domain water gate. Protein, teal cartoon; tunnel surface, orange; water molecules, red spheres.

### Hybrid QM/MM calculations reveal water-mediated catalysis

Wat134 lowers the kinetic barrier for proton-transfer by 20 kcal/mol, enabling a two-step adduct formation: proton transfer from Cys57 to FMN N5, followed by C–S bond formation. Hybrid QM/MM calculations reveal that water-mediated proton transfer via a hydronium-ion transition state faces a barrier of only 15.2 kcal/mol, compared with 35.3 kcal/mol for direct transfer in the absence of the bridging water (Fig. 3b). By Eyring transition-state theory^22^, proton transfer without Wat134 would be ∼10^14^-fold slower (half-life ∼237,000 years versus ∼16 ms with water).

Following proton transfer, the major barrier to adduct formation is reorientation of Cys57 so that its sulphur is positioned over the C4a atom (Fig. 3a,b). If cysteine rotation proceeds in the singlet state immediately after proton transfer, the barrier is prohibitive (∼83 kcal/mol; pathway P-I, Supplementary Fig. 9). In the triplet state with water still bound, it drops to ∼35 kcal/mol (P-II). If water departs after proton transfer, the most favourable pathway emerges with a barrier of only ∼6 kcal/mol (P-III). After cysteine rotation in the triplet state, the reaction crosses to the singlet surface and completes adduct formation without further barrier. These calculations converge on a mechanism in which triplet-state, water-mediated proton transfer is followed by water release, cysteine reorientation, and adduct formation. Time-resolved infrared spectroscopy provides the direct test of this predicted deprotonation step.

### TR-IR spectroscopy resolves cysteine deprotonation kinetics

Time-resolved IR spectroscopy (TR-IR) on well-hydrated CrLOV1 (Extended Data Fig. 6, Supplementary Text) reveals a negative band at 2,568 cm□^1^ together with a red-shifted positive band at 2,542 cm□^1^ in the early nanoseconds (Fig. 5b). The negative band was assigned to the Cys57 thiol^23^, and the positive band to a stronger hydrogen-bonding environment of the S–H group than in the dark state (Fig. 5a). A similar assignment has been reported for the LOV2 domain of Adiantum Phytochrome3^24^. Correlating with the crystallographic data, the hydrogen-bond acceptor is identified as Wat134. Its arrival at Cys57 lowers the pK_a_ and facilitates proton transfer to N5 of FMN.

**Fig. 5:**
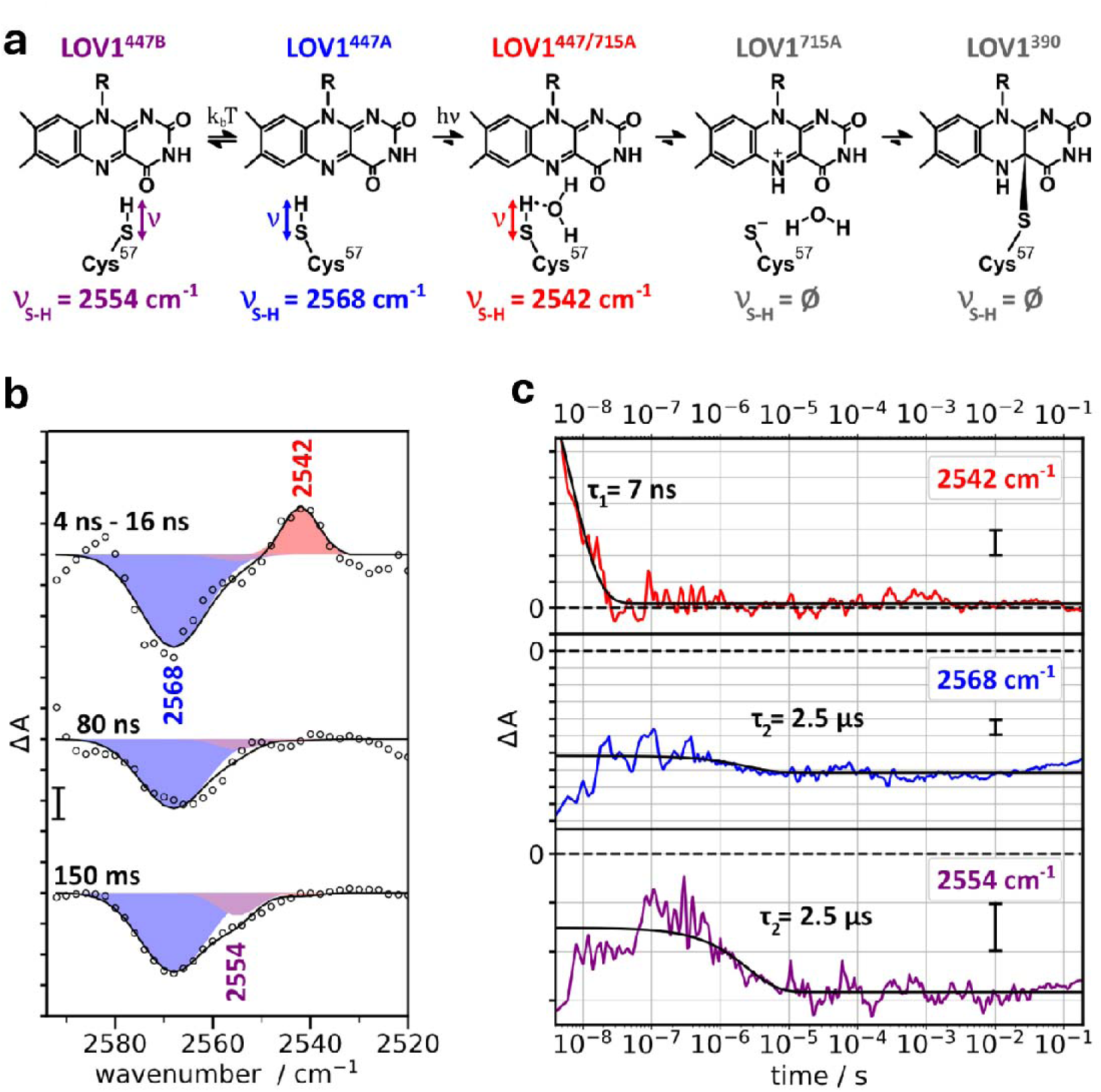
Time-resolved infrared spectroscopy of the Cys57 S–H stretching mode confirms deprotonation at 7 ns, independently corroborating water-mediated proton transfer and resolving two rotamer-specific kinetic pathways. **A.** Proposed reaction model for *Cr*LOV1 photoactivation and assignment of Cys57 S–H stretching bands observed by TR-IR spectroscopy. The 2,568 cm□^1^ band is assigned to the deprotonated or altered-H-bonding thiol; the 2,542 cm□^1^ band to the Cys57 S–H hydrogen-bonded to Wat134; the 2,554 cm□^1^ band to a second Cys57 rotamer species. **B.** Light-minus-dark IR difference spectra recorded with a tunable quantum cascade laser at 4–16 ns (averaged), 80 ns, and 150 ms across the S–H stretching region (2,515–2,590 cm□^1^). Band shapes, Gaussian fits; peak frequencies annotated. Scale bar, 50 µOD. **C.** Kinetic traces at 2,542 cm□^1^, 2,568 cm□^1^, and 2,554 cm□^1^ recorded from 2 ns to 200 ms. The 2,542 cm□^1^ band decays with τ□ = 7 ± 1 ns; the 2,568 cm□^1^ band persists throughout the measured time range, confirming sustained Cys57 deprotonation. The 2,554 cm□^1^ shoulder decays with τ□ = 2.5 ± 1.2 µs, identifying deprotonation of the second Cys57 rotamer at a kinetically distinct timescale. Uncertainties represent standard errors of single-exponential fits. Black traces, exponential fits; scale bars, 20 µ OD.

The initial structural change of the S-H stretch occurs within the response time of the IR detector (τ_Detector_ = 15 ns). The 2,542 cm□^1^ band decays with an apparent τ = 7 ± 1 ns (Fig. 5c), while the negative band at 2,568 cm□^1^ persists (Fig. 5c), indicating the deprotonated state of Cys57 over the entire measured time range after pulsed excitation. Thus, deprotonation of the reactive-cysteine occurs on a timescale similar to intersystem crossing (τ = 3.5 ns), but distinct to adduct formation (τ = 800 ns)^14^ — experimental evidence for temporal separation of these two events, as predicted by our QM/MM calculations (*vide supra*).

The negative shoulder at 2,554 cm□^1^ (Fig. 5b,c τ = 2.5 ± 1.2 µs) indicates the deprotonation of a second cysteine species, assigned to the S–H stretch of Cys57 in the alternative rotamer conformation. The two Cys57 rotamers thus deprotonate at timescales differing by more than two orders of magnitude, resolving the molecular basis for the two adduct-formation rate constants reported by UV/Vis spectroscopy^14^. Adduct formation was independently followed through the FMN C4=O stretching modes (Extended Data Fig. 7, Supplementary Text).

### Cysteine conformational redistribution and covalent bond formation

Covalent thioether bond formation between Cys57 and FMN C4a - the hallmark of LOV photoactivation - occurs on the microsecond timescale. Before bond formation, the Cys57 conformational population redistributes: from 0.52:0.48 (A:B) in the dark state to 0.33:0.67 at 500 ns (+3.6/−4.1 σ; Extended Data Fig. 8), as determined by occupancy refinement. At 1 µs, conformation A converts fully to B, and 45% of conformation B shifts 1.4 Å toward the FMN ring, marking the onset of covalent bond formation (Fig. 3a).

Covalent bond formation drives the C4a carbon from planar sp^2^ to tetrahedral sp^3^ hybridization. This transition initiates at 500 ns, preceding the spectroscopic decay constants of LOV1_715a_ (800 ns) and LOV1_715b_ (4 µs)^14^. At Δ*t* = 1 µs, partial covalent adduct formation is observed at 45% occupancy (Fig. 3a). At Δ*t* = 10 µs, the thioether bond is fully formed (+7.6/−7.9 σ in F_10µs_–F_dark_), and the sp^2^-to-sp^3^ transition of the reactive carbon is completed with characteristic 109° bond angles (Fig. 3a). The remainder of the isoalloxazine ring retains planarity, with an elevation of the pyrimidine ring (+4.6/−3.6 σ at 10 µs).

Upon adduct formation, the nanosecond transient changes reverse: the phosphoribityl tail and Arg58/Arg74 return closer to dark-state positions, and Wat134 disappears entirely by 1 µs (Fig. 3a). These reversals match the QM/MM prediction that N5 protonation precedes bond formation. The incipient C4a shift at 500 ns indicates that bond formation follows protonation almost instantaneously, with A-to-B conformational conversion potentially rate-limiting in a subpopulation.

### Late rearrangements of Gln120 and implications for signal transduction

The Gln120 side chain shifts by 0.2 Å after N5 protonation (+3.2 σ in F^1µs^–F^dark^; Extended Data Fig. 9a), becoming more pronounced at 10 µs (+5.4 σ, Extended Data Fig. 9b). These shifts reorganize the hydrogen-bond network linking Gln120 to FMN and Thr21. The Gln120 amide flips so that its oxygen engages FMN N5 in a new hydrogen bond, disrupting the O4–amide nitrogen interaction and replacing the symmetric 2.8 Å Gln120–Thr21 hydrogen bond with an asymmetric 3.0 Å contact. This weakened N-to-C-terminal interaction defines a structural basis for interdomain signal transduction in full-length phototropin^25^.

## Discussion

A single transiently ordered water molecule is the missing catalyst in LOV domain photoactivation. TR-SFX, supported by QM/MM calculations and time-resolved infrared spectroscopy, reveals the complete structural sequence from FMN photoexcitation in *Cr*LOV1 to covalent thioether bond formation (Fig. 6; Supplementary Movie 1), identifying transient active-site hydration as the critical enabling step. Immediately after light absorption, out-of-plane distortions of the isoalloxazine ring and correlated phosphoribityl tail motion reflect population of the singlet excited state; these distortions relax on the nanosecond timescale as the triplet state forms.

**Fig. 6:**
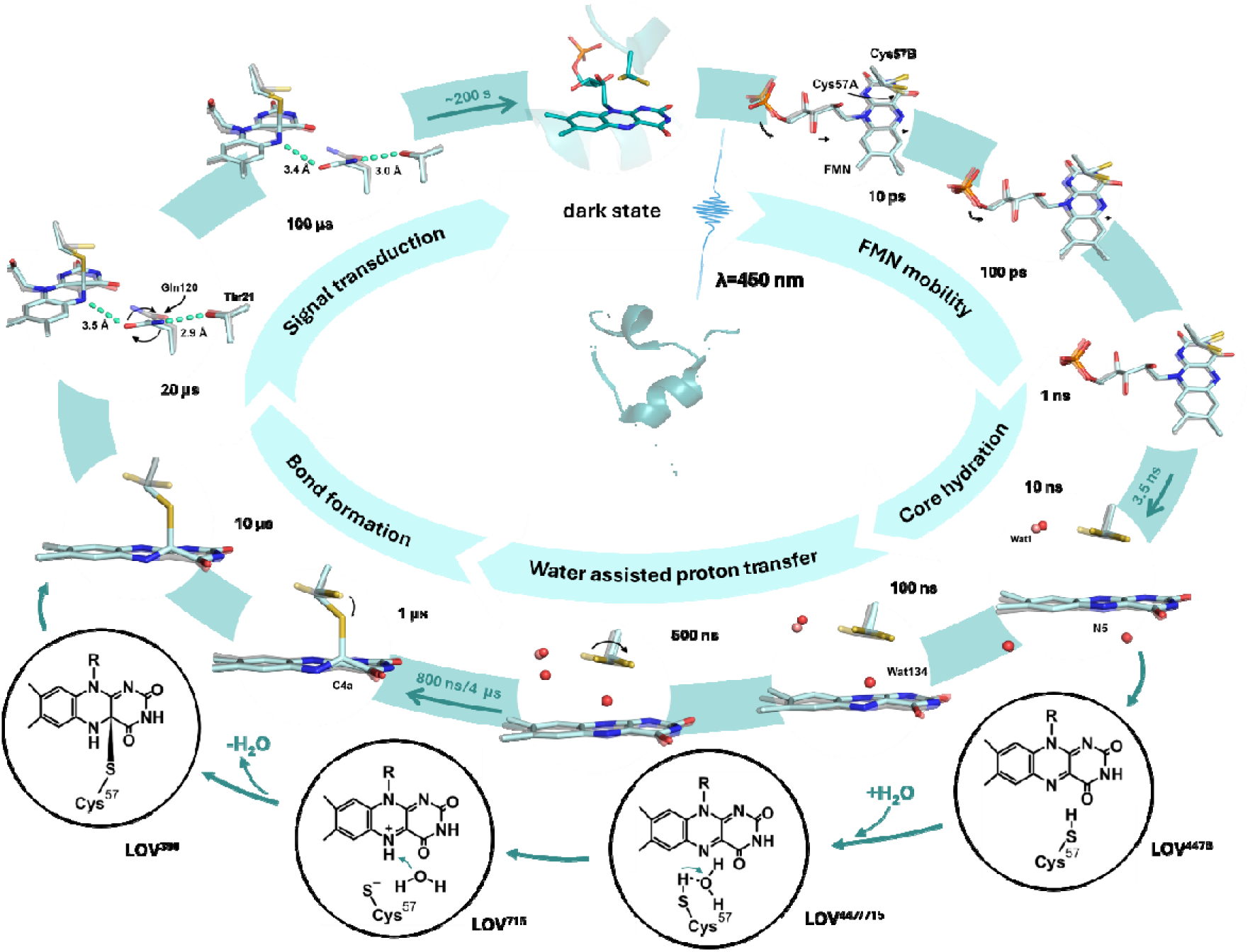
A single transiently ordered water molecule enables an otherwise forbidden proton transfer from Cys57 to FMN N5, completing the *Cr*LOV1 photoactivation cycle from femtosecond excitation to microsecond covalent adduct formation. Radial mechanistic overview of the complete *Cr*LOV1 photocycle. Centre: *Cr*LOV1 secondary structure with bound FMN chromophore. Middle ring: principal mechanistic stages in temporal order (clockwise from top): dark state → blue-light absorption → isoalloxazine ring distortion (ps) → phosphoribityl tail dynamics and global structural expansion (ns) → transient active-site hydration and Wat134-mediated proton transfer from Cys57 to FMN N5 → cysteine conformational redistribution → covalent thioether bond formation → photocycle recovery (∼200 s). Outer ring: representative structural snapshots of the FMN active site at the corresponding TR-SFX time delays, each superimposed on the dark-state reference (grey). Lower arc: chemical diagrams of consecutive FMN states (LOV1_447_ → LOV1_715_ → LOV1_390_) illustrating the bond-forming chemistry and the catalytic role of Wat134 in proton shuttling from the thiol to FMN N5. Quantitative structural and kinetic data for each stage are presented in the corresponding Results sections.

The central mechanistic finding is transient hydration of the dehydrated chromophore pocket. The dark-state pocket is essentially anhydrous near FMN, with only Wat1. Upon photoactivation, several solvent molecules penetrate the protein core in early nanoseconds, and Wat134 appears at a previously unoccupied site between Cys57 and FMN N5 at 100–500 ns — coincident with triplet-state population and preceding covalent bond formation-providing direct structural evidence for transient solvent penetration into the chromophore pocket.

Wat134 lowers the proton-transfer barrier by ∼20 kcal/mol (Fig. 3b), converting a transfer that would otherwise be inaccessible on any biological timescale into a facile one. TR-IR spectroscopy independently confirms stronger hydrogen bonding at the Cys57 thiol and subsequent deprotonation in the nanosecond regime, in agreement with the crystallographic intermediates. A similar but broader band at 2,537 cm□^1^ was reported upon trapping the LOV2 triplet state of Adiantum Phytochrome3 at 77 K^26^. This convergence of three independent methods — crystallography, computation, and vibrational spectroscopy — on the same mechanism firmly establishes water-mediated proton shuttling as the catalytic step.

Following proton transfer, Cys57 redistributes toward the reactive rotamer, C4a rehybridizes from sp^2^ to sp^3^, and the thioether bond forms by 10 µs. The disappearance of Wat134 and the relaxation of all transient features toward the dark-state configuration argue that hydration is catalytic rather than structural, enabling chemistry that cannot proceed in the dehydrated active site, then reversing once its role is fulfilled. The two distinct deprotonation timescales of the Cys57 rotamers, resolved by TR-IR, provide a molecular explanation for the two kinetic components of adduct formation^14^, linking crystallographic structural heterogeneity directly to photocycle kinetics.

Our identification of a solvent access tunnel at the site corresponding to the *As*LOV2 water gate^21^ (Fig. 4) provides the first direct structural evidence for such a gating mechanism in a LOV domain, demonstrating that solvent ingress and egress are tightly coupled to the photocycle. Beyond this catalytic role, water continues to participate in LOV signaling at later stages: in *As*LOV2, recent spectroscopic and computational evidence shows that the subsequent mechanical stage of signal propagation involves expulsion of structured interfacial hydration water, driving Jα helix unfolding^27^.

By directly visualizing both the transient water molecule and the associated protein and chromophore dynamics, this work establishes water-mediated proton shuttling as a key catalytic step in LOV photoactivation. More broadly, dynamic hydration emerges as a catalytic strategy for enabling otherwise forbidden proton-transfer reactions in dehydrated enzyme active sites. Such transiently hydrated catalytic sites are widespread among flavoenzymes, cytochrome P450s, and proton-pumping membrane complexes, where water access to buried reaction centres has long been invoked but here directly captured at atomic resolution. Once the covalent adduct has formed — the step mechanistically resolved here — the structural signal must propagate to effector domains, so water participates in LOV signalling at both the chemical and mechanical levels. Engineering of hydration pathways and the cysteine conformational landscape now offers a rational route to tuning switching kinetics in LOV-based optogenetic tools and flavin-dependent photoswitches.

## Methods

### Expression and purification

The gene encoding the LOV1 domain from *Chlamydomonas reinhardtii* phot protein residues 16-133 were inserted into the expression plasmid pET16b between the restriction sites NdeI and XhoI, allowing the expression of a construct with an N-terminal deca-Histidne (10x Histidine) tag^16^. The protein was expressed in *Escherichia coli* strain BL21 DE3 grown in the auto-inducible media ZYP5052^28^. The cells were grown at 37 °C until OD_600_ ∼1.0 then 17 °C overnight for expression. The cells were harvested by centrifugation at 4,500 xg for 20 minutes at 4 °C and flash frozen in liquid nitrogen until use. The cells were resuspended in lysis buffer (50 mM Tris-HCl pH 8.0, 300 mM NaCl, 1 mM MgSO_4_, 10 mM imidazole, 3 ul DNaseI) and lysed using a high-pressure cell disruptor (Constant Systems Ltd) running at 24 kPSI. The lysate was clarified via centrifugation at 14000 xg for 20 minutes at 4 °C. The protein was purified using and Äkta prime system via nickel affinity chromatography (5ml Histrap HP column, GE Healthcare) and eluted with an imidazole gradient (50 mM Tris-HCl pH 8.0, 150 mM NaCl, 10/500 mM imidazole). The fractions containing the eluted protein were pooled and applied to a gel filtration column for size exclusion chromatography (HiLoad Superdex 75 16/600, GE Healthcare) and buffer exchanged into a 20 mM tris-HCl pH 8.0, 100 mM NaCl buffer. The fractions containing the purified sample were concentrated to 10 mg ml^-1^ for crystallization.

### Crystallization

The crystallization protocol was followed as described in Gotthard *et al*.^16^; in summary: the purification tag was removed from the purified protein via limited proteolysis using trypsin (1:10 ratio of trypsin (0.25 mg ml^-1^) to protein). A condition producing high density microcrystals was identified using crystallization screening, with the best condition containing 10mM sodium cacodylate pH 6.5 and 1.0 mM sodium citrate tribasic dihydrate. The sitting drop vapour diffusion method with 2:1 protein to precipitant ratio produced 10-30 µm crystals, appearing in one day at 20 °C. Crystallization was scaled up and improved crystal size homogeneity by the batch crystallization method with seeding. The crystals grown in the first round of crystallization were crushed with a micro tissue grinder (Elvehjem 0.1 ml) and used as seed stocks. The seeds were mixed with the trypsin-digested protein in a 1:10 ratio and added dropwise into Eppendorf tubes containing the precipitant solution at a ratio of 2:1. Crystals appeared the next day with a homogeneous 20 µm size and sedimented at the bottom of the Eppendorf tube. In order to prepare the larger quantities of crystalline samples required for liquid jet method delivery, a simplified version of this protocol was introduced in which the size exclusion chromatography step was replaced by a simpler and quicker buffer exchange using a HiPrep 26/10 desalting column against final buffer (20 mM Tris-HCl pH8, 100 mM NaCl). In addition, crystal size growth was limited to 5 – 10 µm^3^ by prematurely sedimenting the crystals using centrifugation at 800*g* then reusing the non-crystallized supernatant for several successive cycles of crystallization – sedimentation. This protocol allowed to produce roughly 4,000 mg of crystals for subsequent TR-SFX experiments.

### Experimental setup

#### Sample preparation for TR-SFX at SwissFEL

For TR-SFX data collection at SwissFEL, a viscous jet delivery method was employed using a high-viscosity extruder (HVE)^29^. Microcrystals with an average size of ∼20 µm^3^ were sedimented and resuspended in their crystallization buffer to reach a concentration exceeding 10O crystals ml^−1^. A high-viscous solution of hydroxyethyl cellulose (HEC) (24 % (w/v)) was prepared in a solution containing the protein purification buffer and the crystallization condition in a 1:2 ratio and left for hydration for 24h. The crystal suspension was then mixed in a 1:1 ratio with the viscous HEC media, previously identified as an optimal carrier matrix preserving crystal integrity^16^. The crystal–HEC mixture was prepared by gentle mixing in Hamilton syringes using three-way couplers, yielding a homogeneous, highly concentrated sample suitable for TR-SFX.

TR-SFX data were collected in September 2020 at the ARAMIS-Alvra experimental station of the SwissFEL X-ray free-electron laser. Crystals embedded in the viscous matrix were injected into the X-ray interaction region through a 75 µm nozzle at a stream velocity of 28.1 mm s^−1^, ensuring that each X-ray pulse probed a fresh crystal volume. Diffraction data were recorded at room temperature in a 500 mbar helium atmosphere using a JUNGFRAU 4M detector. Crystals were probed using X-ray pulses at a photon energy of 12.1 keV, a pulse energy of 580 μJ and a pulse duration of 25-30 fs (r.m.s) focused to a spot size of 3 x 5 μm (FWHM) at sample position and a 100 Hz repetition rate.

Light-activated datasets were collected using a femtosecond optical pump laser operating at 475 nm with a repetition rate of 100 Hz. Pump energies down to 2.5 µJ were tested; 5 µJ was the lowest that yielded interpretable light-induced difference density. Thus for the TR-SFX measurements laser pulses had an energy of 5 µJ and a duration of 900 fs, focused to a spot size of 39 × 39 µm^2^ (FWHM) at the sample position, corresponding to a peak fluence of ∼290 mJ cm^−2^ and a peak power density of 322 GW cm^−2^ (evaluated, as for the EuXFEL data, at the centre of the Gaussian profile). Under these conditions, an average of 12.9 photons per FMN chromophore was estimated, not accounting for losses due to scattering by the viscous matrix. The 100 ps structural changes were reproduced in two independent experiments at SwissFEL and the European XFEL, despite different excitation wavelengths (475 versus 420 nm), pulse durations (900 versus ∼30 fs) and sample-delivery methods (Extended Data Fig. 1), confirming that they are robust and not specific to a single excitation condition. Data were acquired in a 4:1 light-to-dark scheme, with four diffraction images collected following laser excitation and one image recorded with the laser off. After sufficient dark data had been collected, data acquisition was continued in a 100% light mode, in which all frames were recorded following optical excitation. All dark images were merged into a single reference dataset and used for Fourier difference map calculations.

#### Sample preparation for TR-SFX at EuXFEL

To leverage the potential of European X-ray Free Electron Laser (EuXFEL), we decided to opt for a liquid jet delivery method (GDVN)^30,31^. Previously optimized crystalline samples were concentrated by centrifugation at 1,000g for 10 min at 293 K. The supernatant was removed and crystals were resuspended in stabilization buffer to obtain a 50% crystal slurry. The suspension was filtered through a 20 µm nylon CellTrics filter (Sysmex) to remove aggregates and large crystals. The crystal slurry concentration was adjusted for optimal jetting at 12%. The filtered crystal suspension was transferred into the jet reservoir, placed on a rocking agitator to prevent sedimentation, and connected to an inline cartridge filter containing a 40 µm frit filter. Microcrystals were delivered into the interaction region using a gas dynamic virtual nozzle (GDVN) with a 75 µm inner diameter^32^. Injection was performed at a flow rate of 37.8 µl min^−1^, producing a focused jet with a diameter of approximately 4.8 µm using helium as the focusing gas. Under these conditions, a jet velocity of 34.6 m s^−1^ was measured.

TR-SFX data were collected in February 2025 at the SPB/SFX instrument of the European XFEL (Schenefeld, Germany)^33,34^ using an experimental setup similar to that described previously^35^. X-ray diffraction data were recorded at a photon energy of 11 keV with an average pulse energy of 1.4 mJ and a pulse duration of approximately 25 fs. The mirror-focused X-ray beam size at the sample position was estimated to be 4.7 × 5.3 µm^2^ (FWHM). Diffraction patterns were collected using the Adaptive Gain Integrating Pixel Detector (AGIPD)^36^. Experimental control and data acquisition were performed using the Karabo framework^37^.

For photoactivation, femtosecond optical laser pulses at 840 nm were frequency-doubled to 420 nm using a 0.5 mm thick beta barium borate (BBO) crystal^38^. The resulting excitation pulses had a duration of ∼30 fs and were focused to an elliptical spot of 28.12 × 78.65 µm^2^ (FWHM). At a pulse energy of 1.25 µJ this corresponds to a peak fluence of 50 mJ cm^−2^, i.e. the fluence at the centre of the Gaussian profile, which is the region sampled by the much smaller X-ray probe. This regime corresponds to an estimated average of 2.5 photons absorbed per FMN chromophore, neglecting scattering and reflection losses at the jet surface. TR-SFX data were collected following a pump–probe scheme similar to that described by Pandey et al. X-ray pulses were delivered in trains of 202 pulses at a 10 Hz repetition rate, with an intra-train repetition rate of 564 kHz. Optical laser excitation was synchronized at 141 kHz, resulting in one laser-excited diffraction image followed by three dark images.

#### Time-resolved infrared spectroscopy

Time-resolved infrared experiments were performed using a home-built setup based on a tunable external-cavity quantum cascade laser (EC-QCL) (Supplementary Fig. 10). To resolve the minute absorption changes of a single S-H vibration with ns time resolution, this instrument was modified from a previous setup^39^ to increase the signal-to-noise ratio. First, a reference detection scheme was installed by splitting the QCL beam using a wedged CaFO window. This transmitted more than 90% of the intensity toward the sample and reflected less than 10% toward the reference detector. After the beam splitter, the transmitted and reflected beams were individually attenuated using wire grid polarizers to avoid detector saturation. The reference detector was matched to the sample detector, and data were recorded using a second oscilloscope channel. Transient absorbance differences were calculated from the ratio of the scaled sample and reference data, which greatly reduced the impact of fluctuations in QCL emission on kinetics. These modifications decreased the noise level of the transient absorption changes of < 5x10-5 in the ns range and to ∼ 10-5 in the µs - ms range.

A 5 ns laser pulse, generated by the 3rd harmonic of a Nd:YAG laser (Quanta-Ray, Spectra Physics) driving an OPO (optical parametric oscillator, OPTA, Darmstadt, Germany), was used for pulsed sample excitation. The emission wavelength was set to 447 nm with an energy density of the laser pulse of 4.5 mJ/cm^2^ (±1.0 mJ/cm^2^) at the sample. The excitation laser artifact was removed by incorporating a shutter alternating between acquiring data with and without exciting the sample. The difference was calculated for each pair of measurements. To probe cysteine dynamics, absorption changes in the S-H stretching vibrational band were sampled across the time range of 2 ns - 188 ms. Kinetic recordings were performed in the frequency range of 2,592–2,482 cm^-1^ in steps of 2 cm^-1^. For experiments in the C=O stretching range, a QCL tunable across 1,800–1,686 cm^-1^ was used. Single kinetic traces were recorded at 1,724 and at 1,711 cm^-1^ with 168 averages collected at each wavenumber. To account for the detector’s decay time (Supplementary Fig. 10), each time trace was corrected using an instrument response correction factor with a time constant of 15 ns. (1/(1-e^(-t/τ))) .

Due to *Cr*LOV1’s long photocycling time (τ = 200 s)^14^, we used a motorized sample wheel to exchange samples. Eight kinetic traces were averaged for each sample at a repetition rate of 2 Hz. The following reasoning justifies this approach: 1) only 10% of the sample molecules are excited per laser pulse, and 2) the photocycle intermediate that populates after 0.5 seconds does not absorb at 447 nm^14^. After eight acquisitions, the sample wheel advanced to the next sample slot, and the procedure was repeated. Once all samples had been measured, the wheel returned to the first sample to record data at the next wavenumber. A waiting time of 450 s was set between wavenumbers to ensure full recovery of the initial dark state before returning to the same sample. The procedure resulted in a total of 288 time traces being recorded for each wavenumber.

From the matrix of recorded time traces at each wavenumber, difference spectra were extracted at specific times after pulsed laser excitation. These time-resolved IR difference spectra were smoothed using a Savitzky-Golay filter with a window length of nine data points and a third-order polynomial (Supplementary Fig 11a). To remove the major contribution of transient heating after pulsed excitation (thermal relaxation), an averaged kinetic trace between 2,530 and 2,500 cm^−1^ was subtracted, as previously described^40^ (Supplementary Fig. 11b). Minor residual heating signals were removed by fitting a first-order polynomial between 2,592 and 2,534 cm^−1^ and subtracting it from the spectrum at each time point as a baseline (Supplementary Fig. 11d). Peak areas were calculated by fitting and integrating Gaussians. The peak positions were fixed at the frequencies of 2,568 cm^-1^, 2,554 cm^-1^, and 2,542 cm^-1^ that were previously identified by the second derivatives of the IR spectra. The time constants of the observed reaction steps were determined by fitting monoexponential functions to the time traces of the frequencies of interest. For the late kinetic event, a global fit was performed simultaneously on the 2,568 cm^-1^ and 2,554 cm^-1^ traces. Uncertainties in the time constants were estimated as three standard deviations (3σ = 99.7% confidence interval), where the standard deviations were calculated as the square roots of the diagonal elements of the covariance matrix returned by the fit.

For experiments within the S-H stretching range, 105 μL of a 200 μM protein sample in buffer solution was pipetted onto a BaFO window in 15 μL increments. After each addition, the sample was allowed to dry for 20 minutes under a controlled anaerobic atmosphere within a glove box. Under aerobic conditions, 10.5 μL of a 50% glycerol/water mixture was placed next to the protein film to allow rehydration via the gas phase. Then, the sample was sealed with a 150 μm spacer, grease, and a second BaFO window. This procedure ensured well-hydrated samples, as confirmed by their FT-IR spectra (Extended Data Fig. 6). A similar sample preparation protocol was applied for IR recordings in the C=O stretching spectral range, but a 100 μm spacer was used to achieve a lower optical density of the sample.

#### Data processing

Serial diffraction images were processed using CrystFEL v0.9^41^. Indexing was performed using the XGANDALF^42^ and MOSFLM^43^ algorithms searching for peaks with a signal to noise ratio of 5.0 and minimum pixel count of 2 and threshold of 1000. The unit cell parameters of the cryogenic temperature structure of CrLOV1 were used with a tolerance of 10 Å for a, b, c dimensions and 2° for α, β and 3° for γ. Peak intensities were integrated using the rings method with indexing radius 3,5,6. Diffraction data corresponding to each of the time delays or the dark data were merged into separate datasets using partialator^44^. During merging of the intensities, a unity model not accounting for partialities was used and the resolution limit was extended by 1-2 nm^-1^. Data collection statistics are presented in table S1.

#### Extra-Xwiz pipeline for data processing at EuXFEL

Data processing for 100ps collected at EuXFEL was performed using EXtra-Xwiz^45^ with CrystFEL 0.11.1^41,46^. Peak finding was carried out using the peakfinder8 algorithm^47^, and integration was performed using XGANDALF^42^ with multi-crystal indexing enabled. The peakfinder parameters were set as follows: peak threshold = 3500, peak signal-to-noise ratio (SNR) = 4.5, and minimum number of peaks = 15. Ring integration radii of 3, 5, and 7 pixels were applied. Detector geometry was refined iteratively using geoptimiser^48^, detector-shift, and millipede^49^ over several cycles.

#### Difference Fourier electron density maps and extrapolated structure factors calculations

Fourier difference electron density maps were calculated using *phenix.fobs_minus_fobs_map* with the use of the multiscaling method, a high resolution cutoff corresponding to the highest resolution of the light data and an *F*_obs_ cutoff of 3.0 σ using the phases from the dark state reference. Light-induced structural changes occurring at the different timepoints were modeled using extrapolated structure factors calculated as reported previously^16,50^ with the following formula *F*_ext_ = [(*F*_obs_^light^ – *F*_obs_^dark^) / α] + *F*_obs_^dark^ where α corresponds to the activated fraction, *i*.*e* account for the contribution of non-activated molecules that are mixed with triggered ones following light illumination of the crystal sample. First extrapolated electron density maps were calculated using the dark state model at decreasing activation levels until the dark state features appeared less predominant while monitoring the overall quality of the maps. A conservative resolution cutoff accounting for error propagation in extrapolated structure factors calculation was applied such as R_free_ was below 50% in the highest resolution shell. Respective activation levels and refinement statistics are reported in Extended Data Table 1.

#### Model building and refinement

Structures were solved using molecular replacement with the program Phaser^51^ using the structure coordinates of the LOV1 domain from *Chlamydomonas reinhardtii* phot (PDB code 1N9L)^15^ as a search model. Iterative cycles of model building and refinement was carried out against the 2F_extrap_-F_c_ electron density maps using the graphic software Coot^52^ and PHENIX software suite^53^. In order to accurately model the high energy intermediate states of the chromophore, relaxed torsion angle restraints were generated, and the modified cif restraints files were used during the structure refinement. The covalent adduct was modelled by supplementing the refinement input files with a parameter file, describing the geometry of the covalent bond between the Cys57 sulphur and FMN C4a atom. Structure analysis and visualization was prepared using the Open-Source PyMOL (http://pymol.org/pymol, The PyMOL Molecular Graphics System, version 2.6, Schrodinger, LLC.). Coordinates and structure factors have been deposited to the Worldwide Protein Data Bank under the access codes 30NO, 30ML, 30MI, 30MK, 30QF, 30MM, 30MJ, 30ST, 30SE, 30SU, 30SV, 30ZY.

#### QM/MM methods

The crystal structure obtained at 500 ns delay was used as the starting point for hybrid quantum mechanical/molecular mechanics (QM/MM) simulations. Based on spectroscopic evidence for an intersystem crossing between the singlet to the triplet state on the early nanosecond timescale, geometry optimization of the flavin mononucleotide (FMN) was performed in the triplet excited state. The chromophore was initially modelled in its oxidized form using force-field parameters from Schneider *et al.*^54^, and protonation states of the protein residues were assigned using the *tleap* module of the AMBER software package^55^.

During geometry optimization, the QM region comprised the FMN chromophore, Cys57, and a nearby water molecule (Wat134). FMN was truncated between the C1′ and C2′ atoms, and Cys57 between the Cα and Cβ atoms; hydrogen link atoms were introduced at the QM/MM boundaries. A spherical region with a radius of 5 Å from each atom of the chromophore was treated as flexible during optimization, while the remainder of the system was held fixed.

To investigate the proton-transfer energetics, the reduced form of FMN was first optimized in the triplet state with Cys57 deprotonated, thereby mimicking proton transfer to the chromophore. The minimum-energy pathway and transition state for proton transfer from Cys57 to FMN were then determined using the nudged elastic band (NEB) method^56^. Additionally, calculations were also performed with the reactive water molecule removed from the active site to assess its role in the process. Approximate proton-transfer rate constants were estimated using the Eyring transition-state theory^22^:

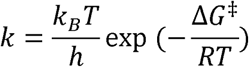

where k is the rate constant, ΔG^‡^ is the Gibbs free energy of activation (reaction barrier), k_B_ is the Boltzmann constant, T is the temperature (300 K), and ù is the Planck constant.

Adduct formation was investigated using an approach analogous to that described above. Geometry optimizations were performed for the adduct in both singlet and triplet states; however, stable adduct formation was observed only in the singlet state. Consequently, subsequent NEB calculations were carried out in the singlet manifold, starting from the proton-transferred, reduced FMN. Prior to the NEB calculations, the reduced FMN was reoptimized in the singlet state to ensure a consistent reference geometry for probing the adduct-formation barrier.

All triplet-state calculations employed the restricted open-shell CAM-B3LYP functional^57^ with dispersion correction^58^ in conjunction with the cc-pVDZ basis set^59^. Singlet-state calculations were carried out using the corresponding restricted closed-shell formalism. Protein residues were described using the AMBER ff14SB force field^60^, and water molecules were modelled with the TIP3P water model^61^. All geometry optimizations and NEB calculations were performed using TeraChem^62,63^ interfaced with ChemShell^64,65^.

## Data and materials availability

Coordinates and structure factors have been deposited to the Worldwide Protein Data Bank under the access codes 30NO, 30ML, 30MI, 30MK, 30QF, 30MM, 30MJ, 30ST, 30SE, 30SU, 30SV, 30ZY.

## Acknowledgments

We acknowledge Alvra Experimental Station of SwissFEL (No. 20200791) and the SPB/SFX Instrument of European XFEL (No. 2645 and 7955) for provision of X-ray free-electron laser beamtime and would like to thank the staff for their assistance. We would like to acknowledge Thomas Dietze for his technical assistance. We acknowledge P11 beamline of PETRA III operated by DESY Photon Science (No. R-20240820 EC) and X06SA beamline at the Swiss Light Source (No. 20191892) and would like to thank the staff for their assistance. We acknowledge the Macromolecular Crystallisation Facility at the Paul Scherrer Institute, Protein Crystallization Center University Zurich, Malopolska Center of Biotechnology Structural Biology Core Facility for access and support.

## Author contributions

P.No. designed and coordinated the experiment. I.S. coordinated quantum mechanical calculations. J.H. coordinated time-resolved spectroscopy. G.G., B.O., A.K., M.P., A.S., A.A, R.K., expressed, purified and crystallized *Cr*LOV1. A.K., A.S., R.K., M.P., A.A., J.V, S.M., A.F., P.S., K.D., H.H., J.Sc. and G.G. secured constant supply of sample during the SFX beamtimes. G.G., S.M., A.K. and A.S. optimized crystal injection. P.J.M.J., D.J., K.N., D.G., C.C., C.B., C.M., F.D., T.S., T.P., T.D., J.E., F.T., C.K., R.L., J.B. and R.B. prepared and operated endstations, including the laser systems. G.G., S.M., B.O., D.O., K.N. M.K.S., F.H.M.K., O.T., E.S., D.M., D.Z. and R. dW. performed data processing during beamtimes. M.Wr., B.O., and A.K. recorded progress during data collection. G.F.X.S. and J.St supported crystallographic applications at SwissFEL and contributed to discussions. G.G. optimized data processing. G.G, B.O and P.No refined and interpreted structures. P.Na. and I.S. performed and interpreted QM/MM calculations. M.G.-V., P.L. and J.H. performed and interpreted the time-resolved spectroscopic experiments. B.O. and P.No. wrote the manuscript with direct contributions from G.G., P.Na. M.G.-V., M.Wi., J.H. and I.S., with further contributions from most of the other co-authors. All authors read and approved the manuscript.

## Competing interests

The authors declare no competing interests.

* Correspondence and requests for materials should be addressed to J.H., I.S. or P.N.

## Additional information

Supplementary information is available for this paper containing Supplementary Text, and Supplementary Fig. 1-11.

---

**Extended Data Fig. 1:**
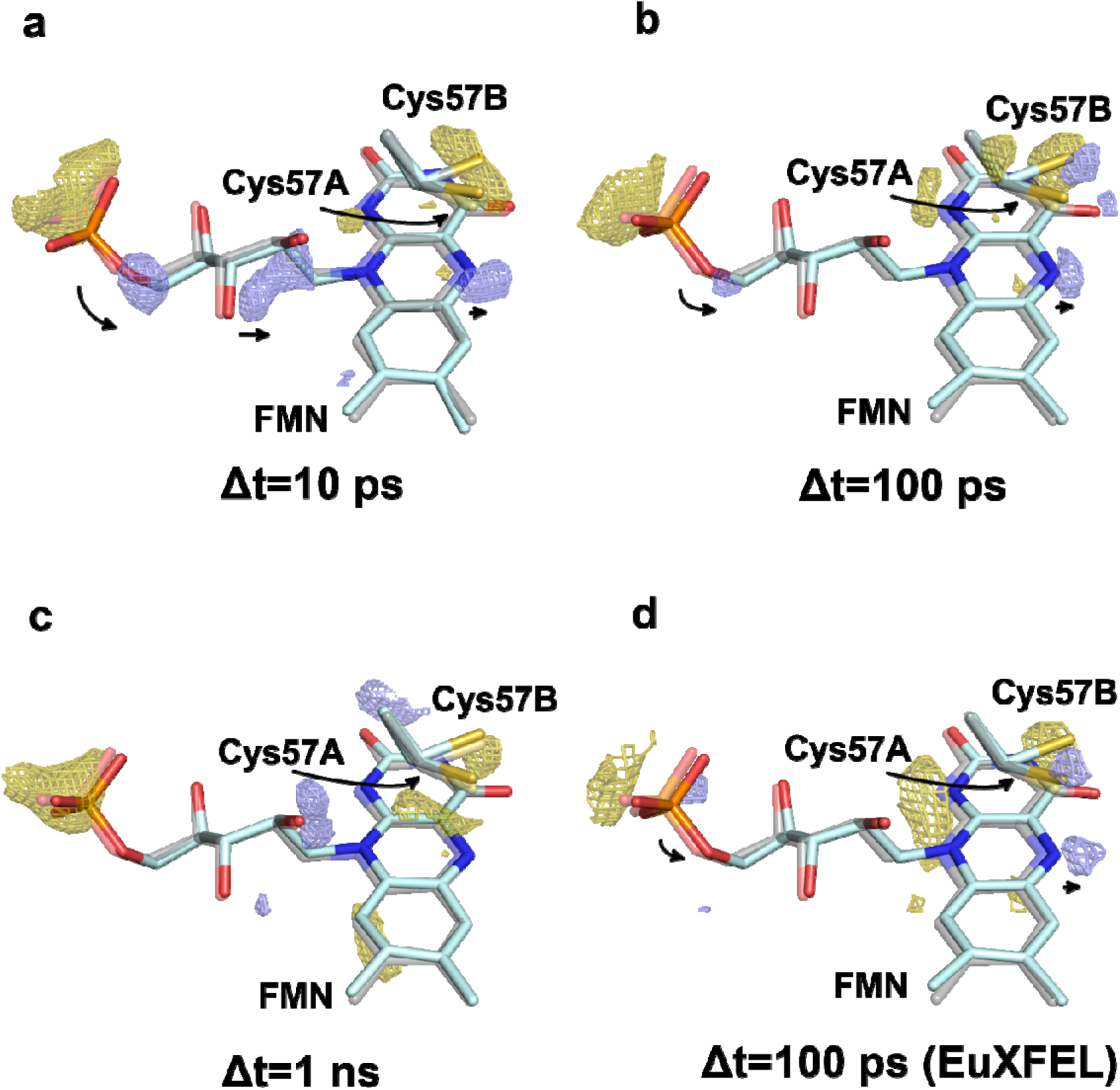
The contraction of FMN phosphate tail at the early timepoints of the photocycle. **A)** The strong negative density at the phosphate group is accompanied by positive peaks along the phosphate tail at the Δt=10 ps intermediate. **B)** The positive density along the tail has disappeared with strong features present at the phosphate group and the N5 atom at Δt=100 ps. **C)** At Δt=1 ns post photoexcitation the negative density is still present at the phosphate group, with the remaining features along the tail and N5 group reduced. **D)** Data collected at the Δt=100 ps time delay from the European XFEL facility. The observed difference density resembles closely the data from the SwissFEL source. The F_obs_^light^ - F_obs_^dark^ difference Fourier electron density map contoured at ±3.0 σ, with positive density represented by blue, and negative density by gold coloured mesh. The excited state chromophore is coloured light blue, overlaid with gray transparent sticks of the resting state structure.

**Extended Data Fig. 2:**
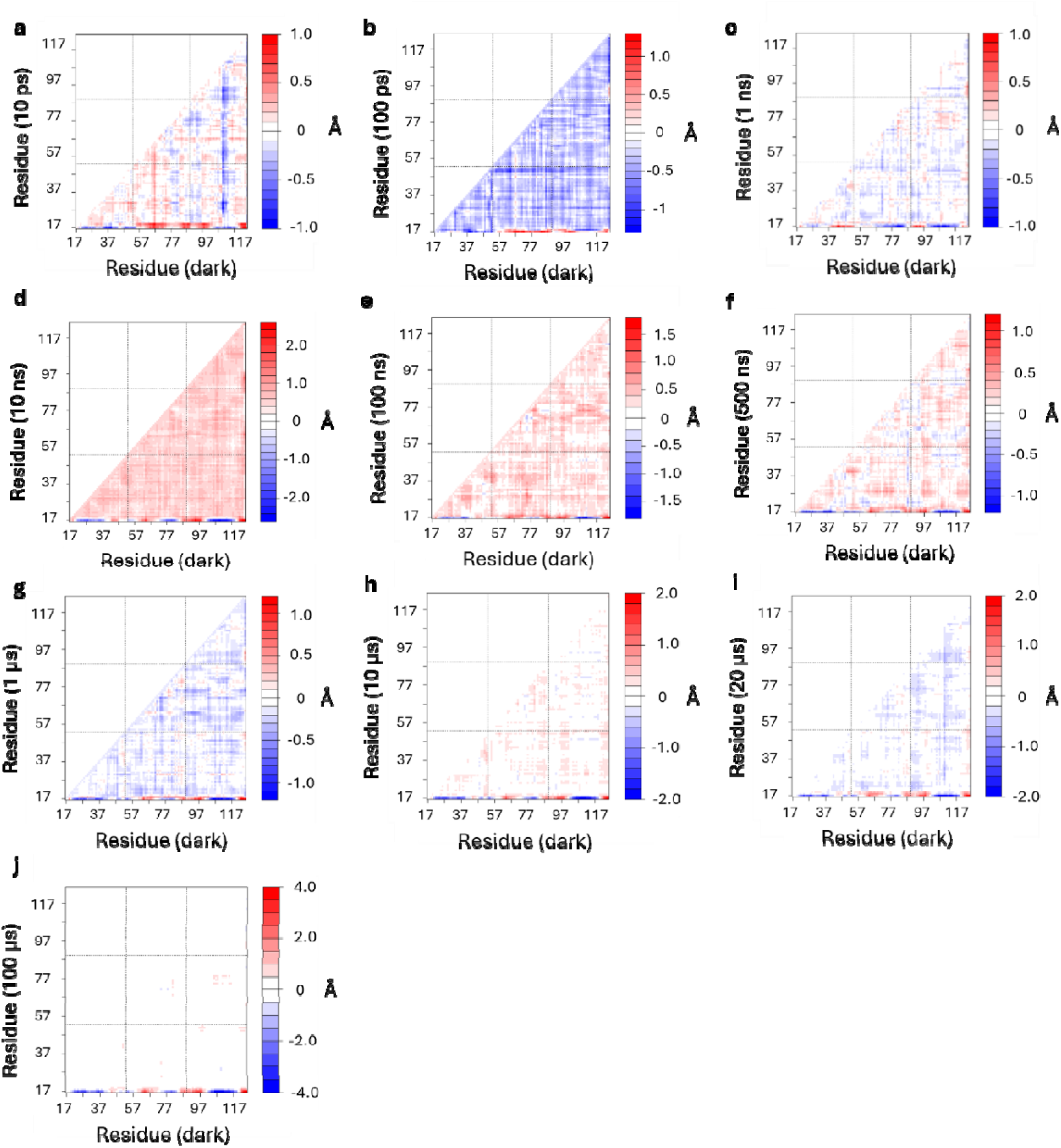
Cα distance matrices of the probed timepoints against the dark state structure. The timepoints representing the singlet excited state show an overall contraction of the protein structure (A-C; 10 ps, 100 ps, 1 ns). This contraction changes to an overall expansion at the triplet excited state timepoints (D-F; 10 ns, 100 ns, 500 ns), coinciding with the observed hydration of the chromophore environment. Once the covalent bond formation occurs, the protein relaxes back to close to the dark state positions of the Cα atoms (G-J; 1 µs, 10 µs, 20 µs, 100 µs).

**Extended Data Fig. 3:**
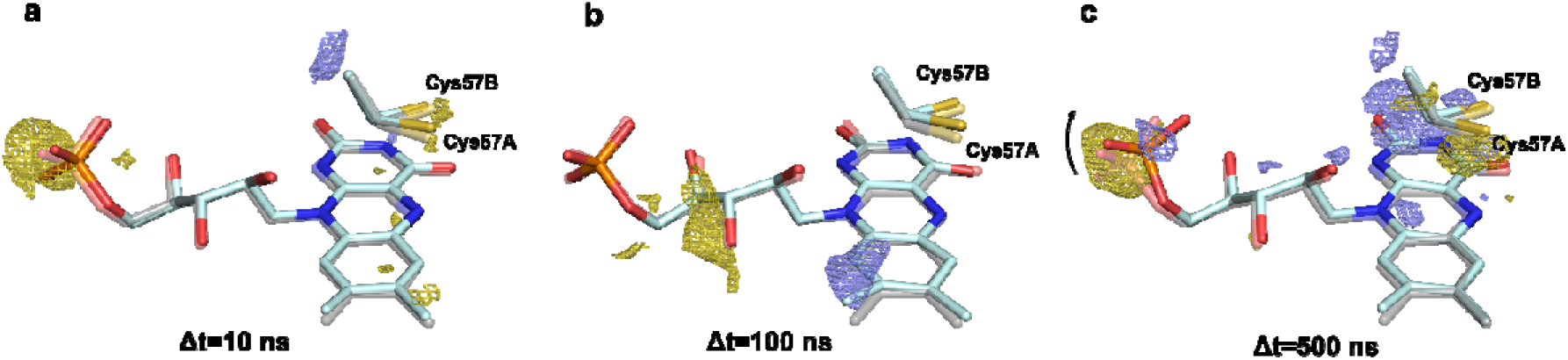
Disorder at the FMN phosphate group in the nanosecond timescale. **A)** The contraction of the phosphate tail relaxes to the resting state position at the Δt=10ns timepoint. **B)** At the Δt=100 ns intermediate, the phosphate group shows no displacement compared to the dark resting state position. **C)** The phosphate group shifts upwards when compared to the dark state position at the Δt=500 ns timepoint, the last timepoint prior to the covalent bond formation. The F_obs_^light^-F_obs_^dark^ difference Fourier electron density map contoured at ±3.0 σ, with positive density represented by blue, and negative density by gold coloured mesh. The excited state chromophore is coloured light blue, overlaid with gray transparent sticks of the resting state structure. Alternate conformations of the reactive Cys57 are labelled as A/B.

**Extended Data Fig. 4:**
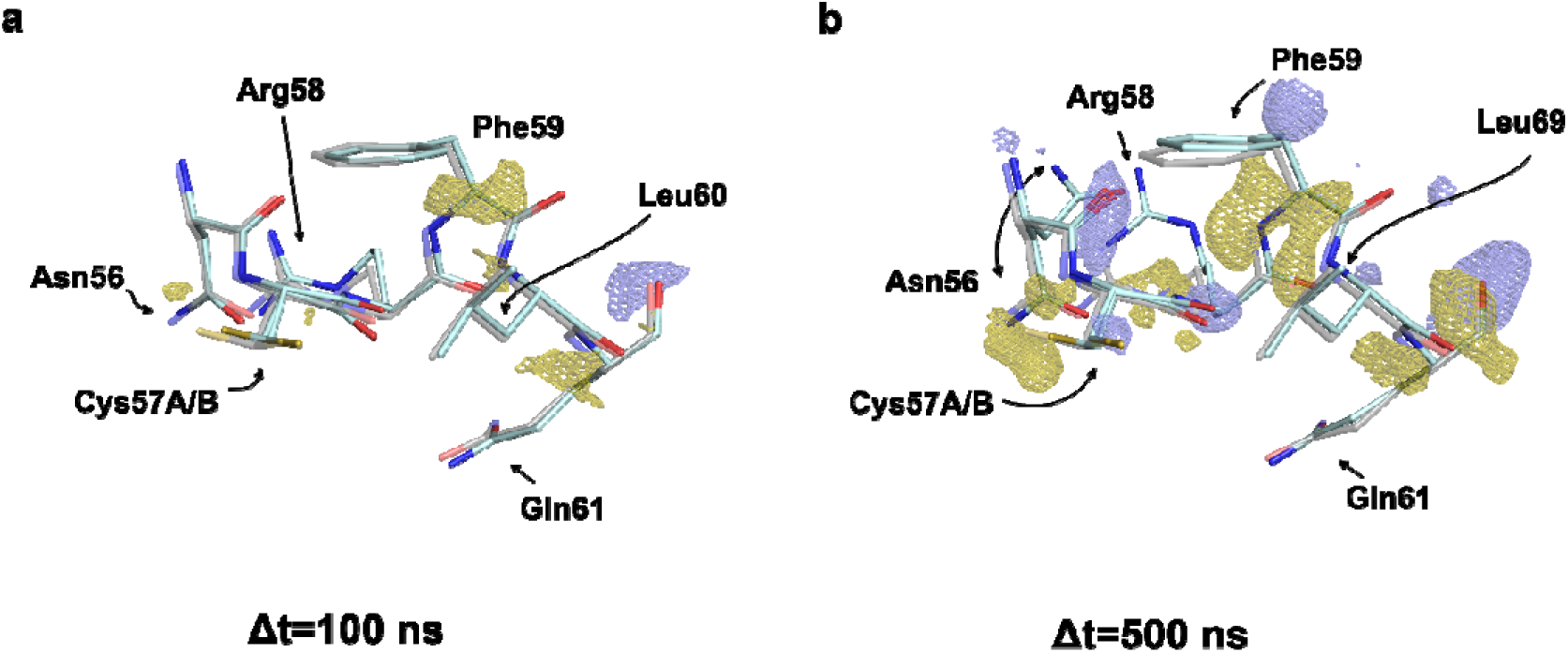
The increased disorder of phosphate tail and surface Arg58 and Arg74 is accompanied by shift in the main chain of LOV in the region of reactive Cys57 for residues Asn56-Gln61. A) At the 100 ns intermediate the main chain of the residues around the reactive cysteine resemble the resting state closely, with few F_100ns_-F_dark_ difference density peaks observed. B) At the subsequent probed timepoint, Δt=500 ns, major shifts are observed accompanied by multiple large F_500ns_-F_dark_ difference density pairs. Besides the main chain shift, Asn56 also undergoes a partial shift, weakening the H-bond network of the phosphoribityl tail stability. The F_light_ – F_dark_ difference Fourier electron density map contoured at ±3.0 σ, with positive density represented by blue, and negative density by gold coloured mesh, displayed around the depicted amino acid residues only. The excited state chromophore is coloured light blue, overlaid with gray transparent sticks of the resting state structure.

**Extended Data Fig. 5:**
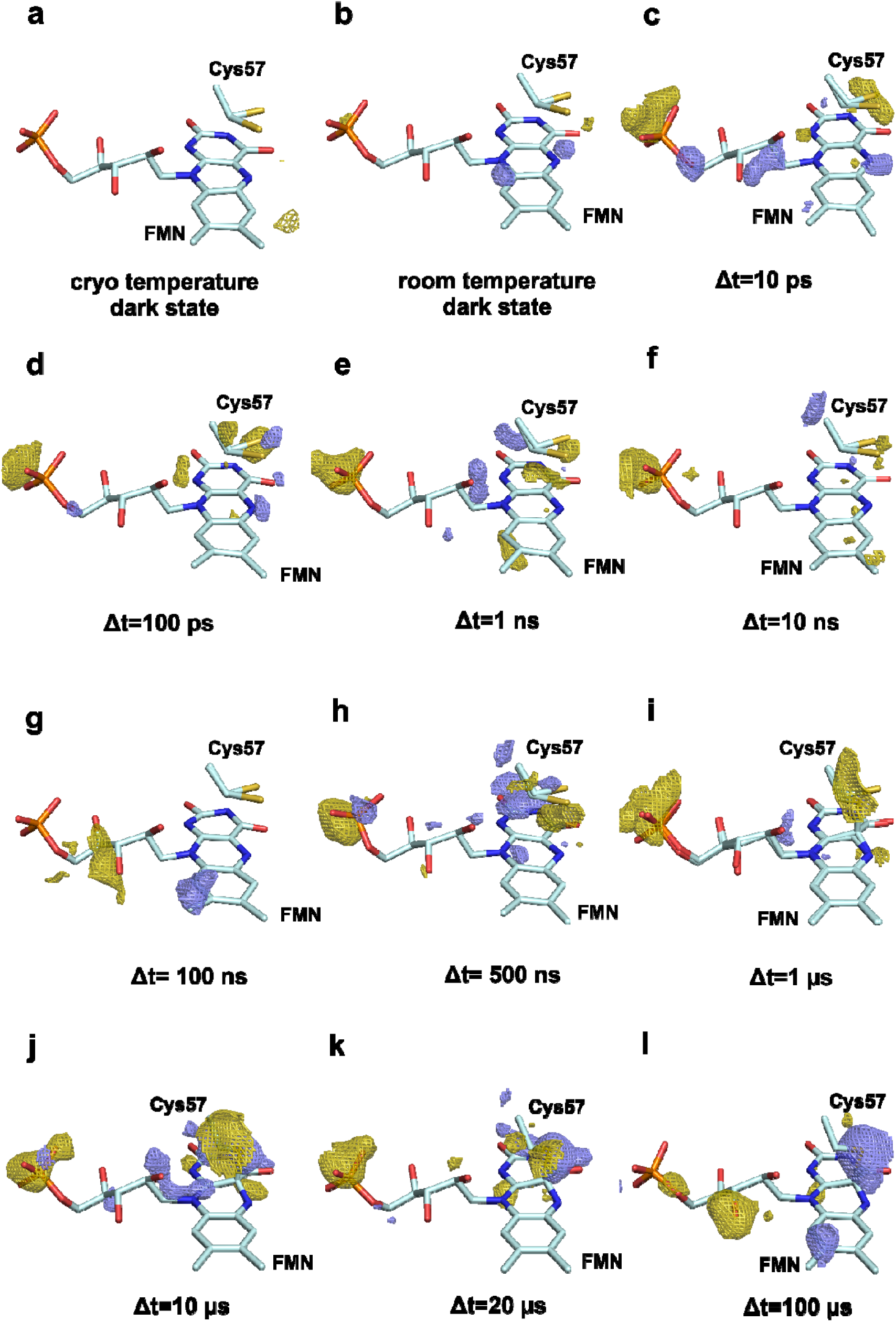
Light activation results in increased disorder at the phosphate tail in all probed timepoints. **A)** The dark state structure captured at cryo temperature (PDB: 1N9L)^15^ shows no difference density near the FMN chromophore. **B)** The room temperature resting state structure captured at the SwissFEL Alvra endstation. An increased mobility is visible compared to the cryo temperature, signified by a -3.3 σ negative density at the phosphate group. **C-L)** Increased mobility of the bulk solvent facing phosphate group is striking with greatly increased negative electron density in the F_obs_^light^-F_obs_^dark^ maps, ranging from -4.4 to -6.9 σ negative densities. The most significant difference density, accompanied by a positive difference density peak (+3.9 σ) is observed at the 500 ns intermediate **(H)**, coinciding with the largest displacement seen in the coordinating arginine residues.

**Extended Data Fig. 6:**
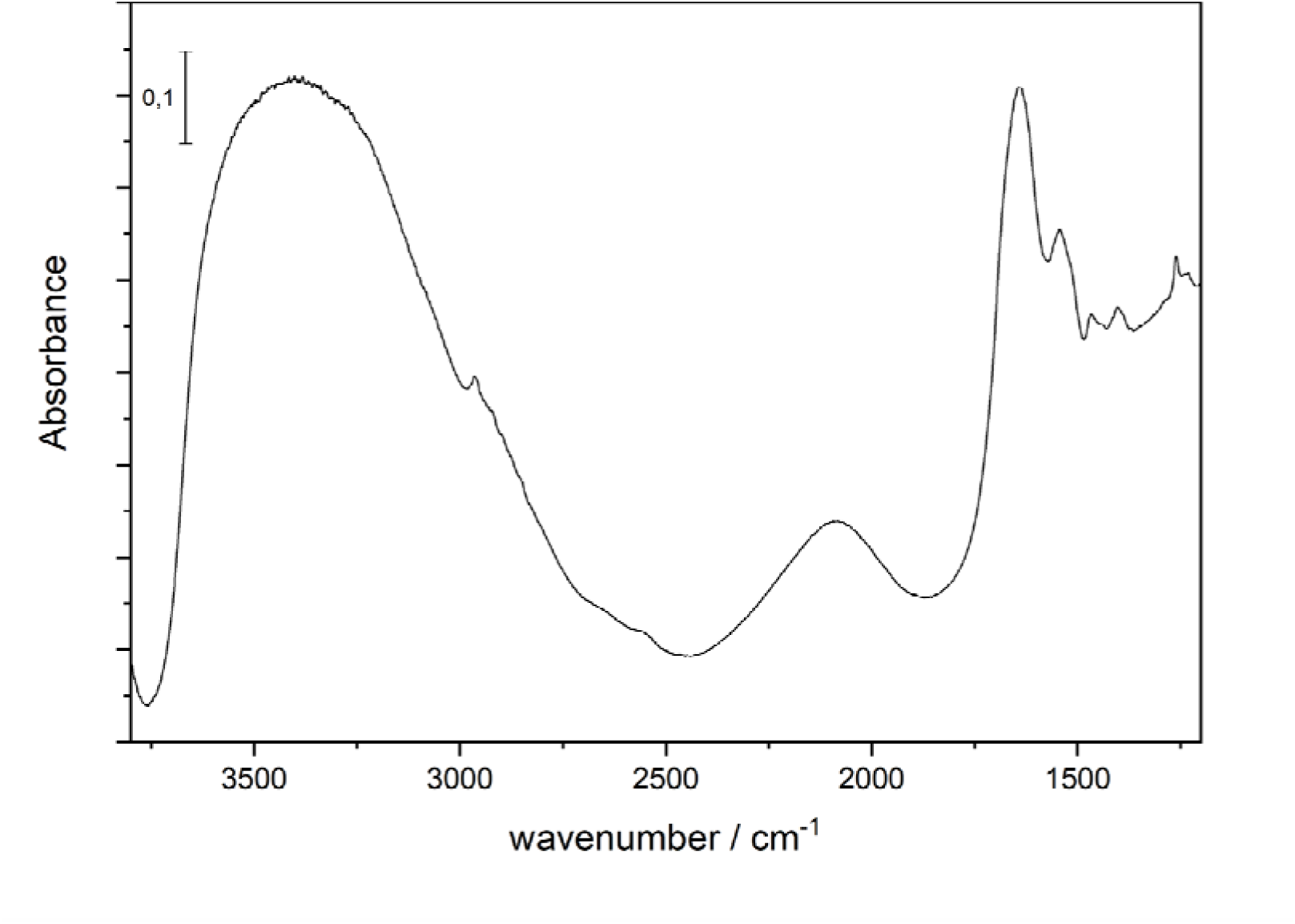
Samples used for time-resolved infrared spectroscopy were maintained over well-hydrated conditions. Absorbance FT-IR spectrum of 45 μL *Cr*LOV1 rehydrated with 10.5 μL 50%(w/w) glycerol/water mixture, and 150 μm spacer thickness.

**Extended Data Fig. 7:**
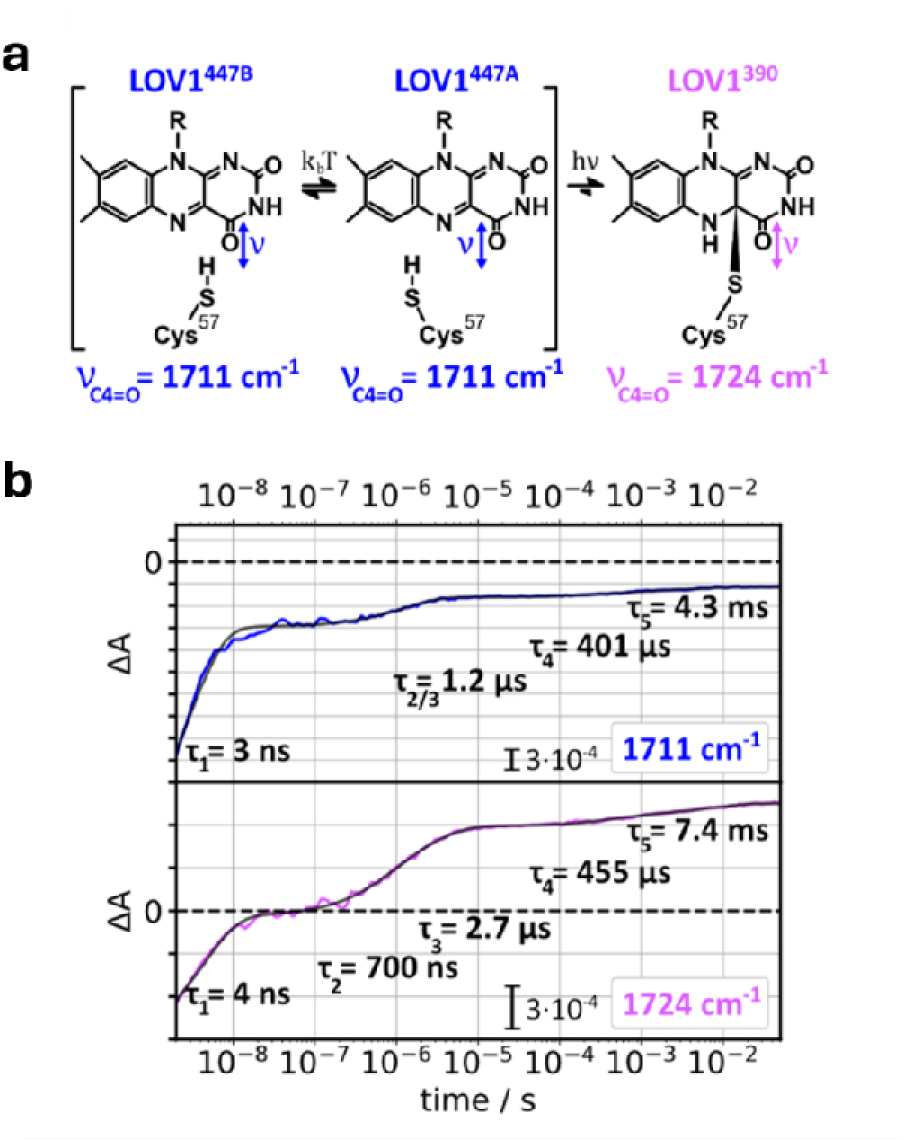
Time-resolved IR difference spectroscopy on CrLOV1 of C4=O stretching of the FMN using a tunable quantum cascade laser. **A)** C4=O stretching modes of the FMN followed in time^16^. **B)** Kinetic traces recorded at 1711, and 1724 cm^-1^ (in colour) over the time range from 2 ns – 200 ms. The black traces are multi-exponential fits with the time constants as shown. (See Materials & Methods for details of the experiment.)

**Extended Data Fig. 8:**
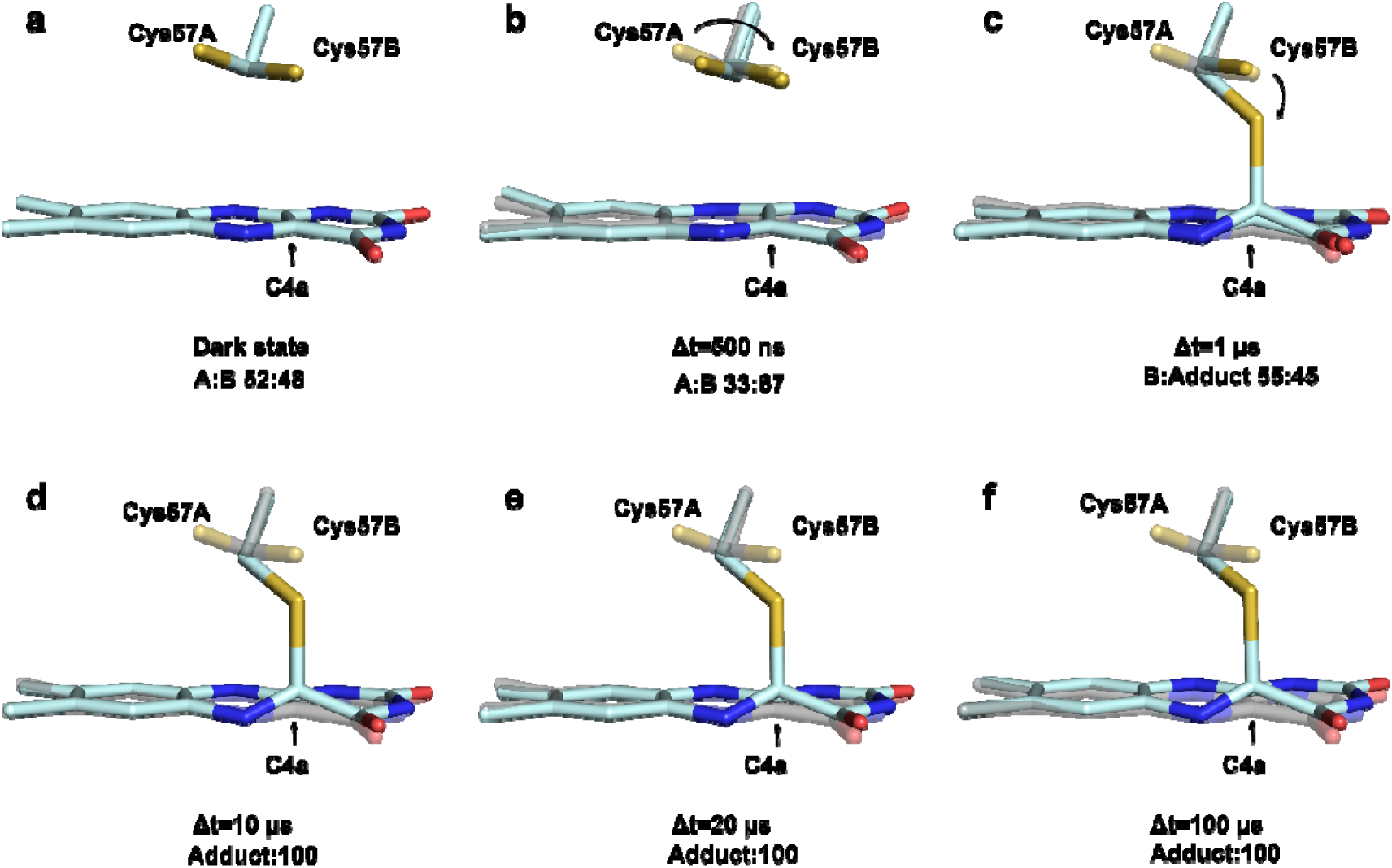
Formation of the covalent bond at the early µs timepoints post photoexcitation. **A)** Dark resting state structure consists of a planar FMN ring structure, even cysteine conformation occupancy. **B)** At the Δt=500 ns timepoint, a shift in the cysteine conformations is observed. **C)** 1 µs post photoexcitation Cys57A conformation disappears, and a partial occupancy bond forms between the protein and the chromophore. **D)-F)** At the 10 µs timepoint/20 µs timepoint/100 µs intermediates, the reactive cysteine participates in the bond, with both conformations from the resting state disappearing. The structure of the reaction intermediates are shown in pale cyan sticks, overlaid with the dark resting state structure in grey transparent sticks. The reactive cysteine sidechains are shown in identical representation, with alternate conformations labelled as A/B.

**Extended Data Fig. 9:**
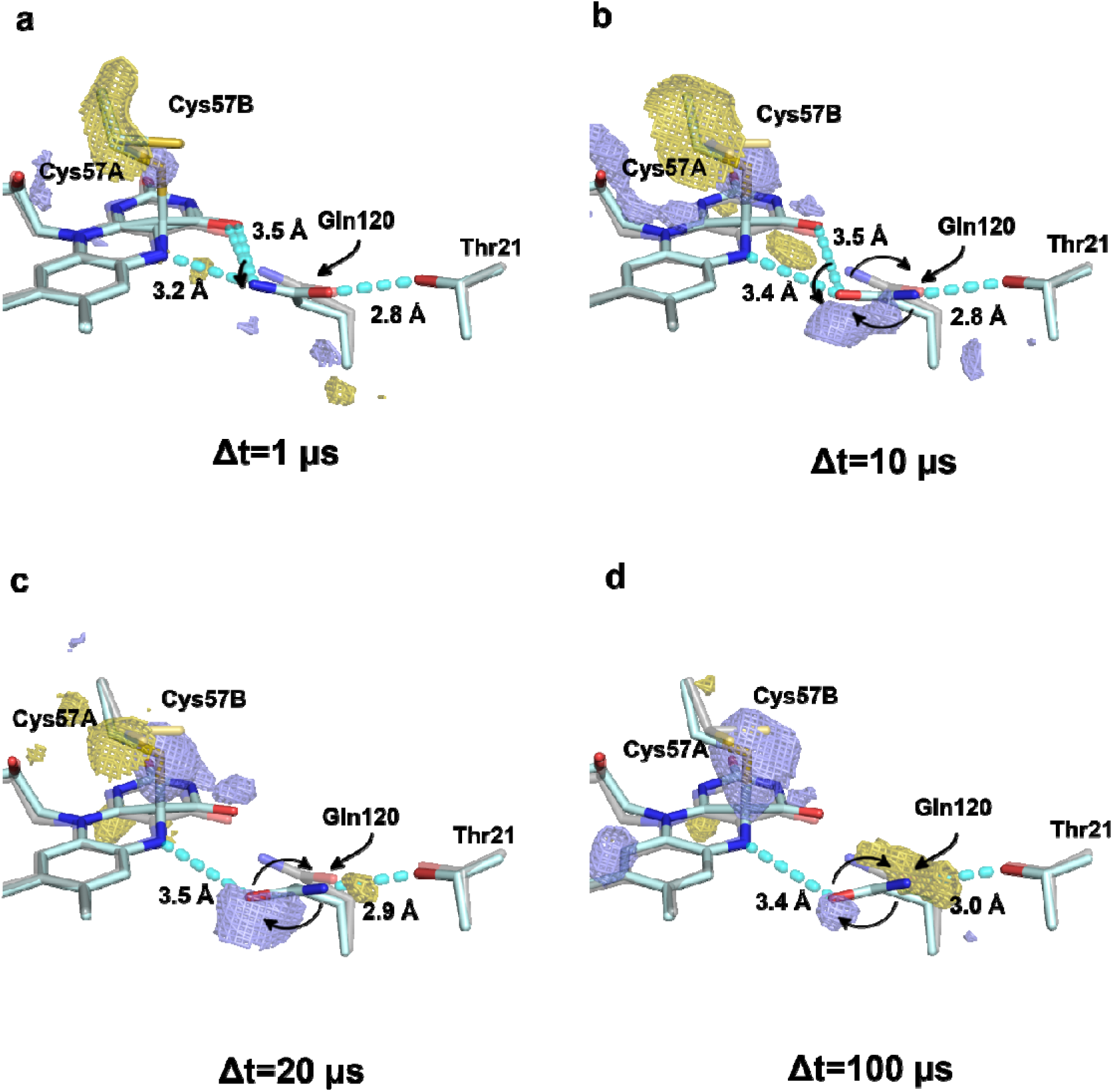
Flip of the Gln120 sidechain and rearrangement of the hydrogen bond at the µs timescale. **A)** At Δt=1 µs post photoexcitation Gln120 shifts slightly from the dark resting state position. **B)** Upon formation of the covalent bond at the 10 µs timepoint, the repositioning of the sidechain is more pronounced, with a large positive difference density peak observed. **C)** Subsequently, the Gln120 sidechain flips, rearranging the H-bond network it forms with the chromophore (Δt=20 µs). **D)** At the Δt=100 µs timepoint the flip is indicated with a stronger negative difference density at the glutamine resting state position. The structure of the reaction intermediates are shown in pale cyan sticks, overlaid with the dark resting state structure in grey transparent sticks. The reactive cysteine and Gln120 sidechains are shown in identical representation. The hydrogen bonds are shown as cyan dashed lines. The F_obs_^light^-F_obs_^dark^ difference Fourier electron density map contoured at ±3.0 σ, with positive density shown as blue, and negative as gold coloured mesh.

**Extended Data Table 1.**
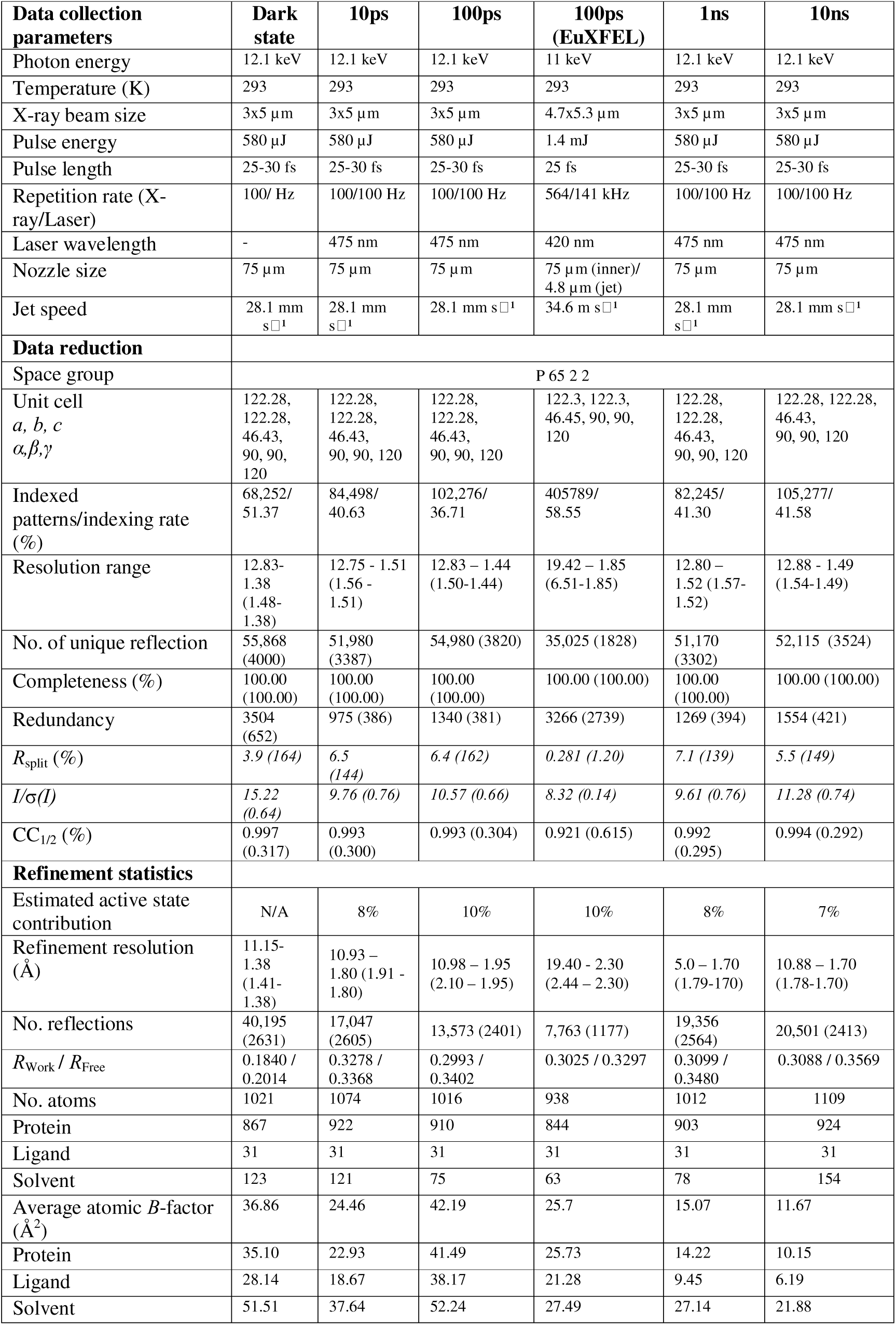
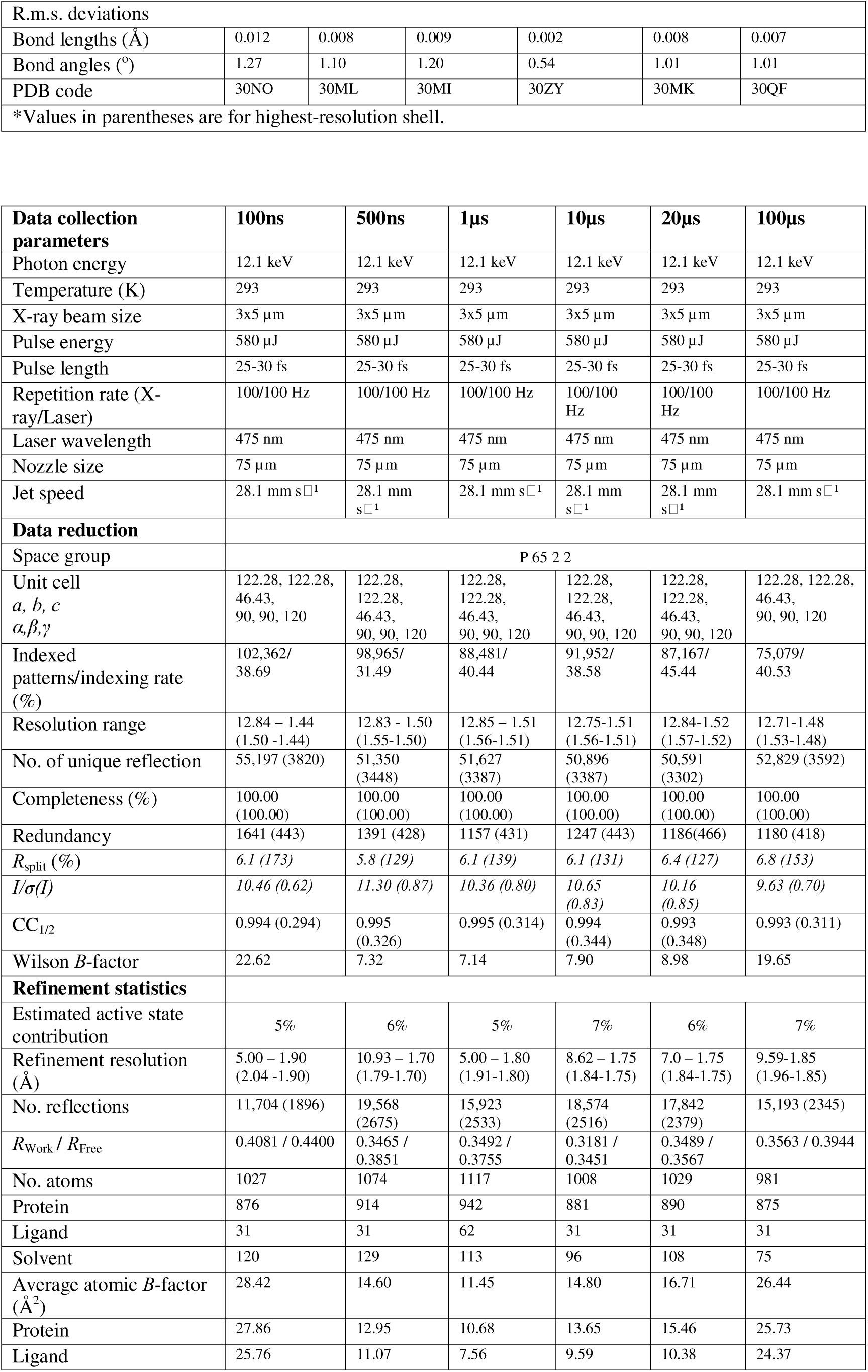
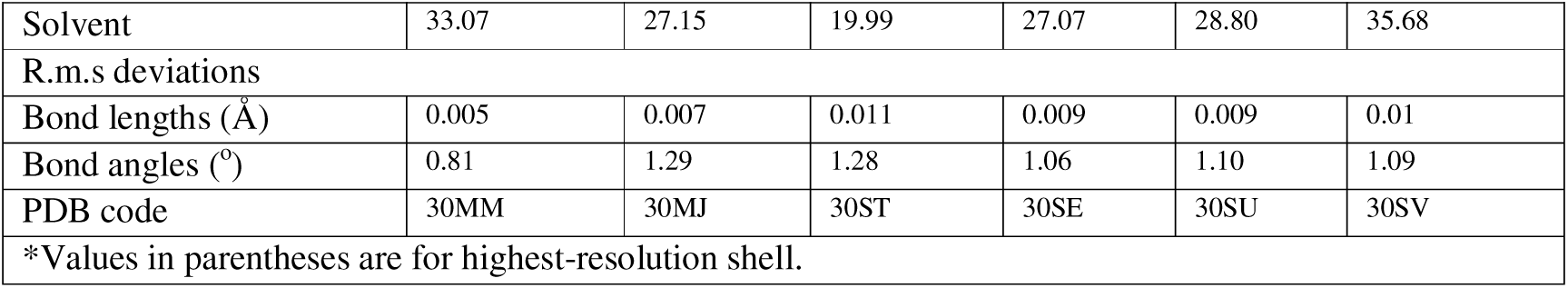
TR-SFX data collection, data reduction and refinement statistics.

| **Data collection parameters** | **Dark state** | **10ps** | **100ps** | **100ps (EuXFEL)** | **1ns** | **10ns** |
| --- | --- | --- | --- | --- | --- | --- |
| Photon energy | 12.1 keV | 12.1 keV | 12.1 keV | 11 keV | 12.1 keV | 12.1 keV |
| Temperature (K) | 293 | 293 | 293 | 293 | 293 | 293 |
| X-ray beam size | 3x5 $\mu\text{m}$ | 3x5 $\mu\text{m}$ | 3x5 $\mu\text{m}$ | 4.7x5.3 $\mu\text{m}$ | 3x5 $\mu\text{m}$ | 3x5 $\mu\text{m}$ |
| Pulse energy | 580 $\mu\text{J}$ | 580 $\mu\text{J}$ | 580 $\mu\text{J}$ | 1.4 mJ | 580 $\mu\text{J}$ | 580 $\mu\text{J}$ |
| Pulse length | 25-30 fs | 25-30 fs | 25-30 fs | 25 fs | 25-30 fs | 25-30 fs |
| Repetition rate (X-ray/Laser) | 100/ Hz | 100/100 Hz | 100/100 Hz | 564/141 kHz | 100/100 Hz | 100/100 Hz |
| Laser wavelength | - | 475 nm | 475 nm | 420 nm | 475 nm | 475 nm |
| Nozzle size | 75 $\mu\text{m}$ | 75 $\mu\text{m}$ | 75 $\mu\text{m}$ | 75 $\mu\text{m}$ (inner)/<br>4.8 $\mu\text{m}$ (jet) | 75 $\mu\text{m}$ | 75 $\mu\text{m}$ |
| Jet speed | 28.1 mm $\text{s}^{-1}$ | 28.1 mm $\text{s}^{-1}$ | 28.1 mm $\text{s}^{-1}$ | 34.6 m $\text{s}^{-1}$ | 28.1 mm $\text{s}^{-1}$ | 28.1 mm $\text{s}^{-1}$ |
| **Data reduction** |  |  |  |  |  |  |
| Space group | P 65 2 2 |  |  |  |  |  |
| Unit cell<br>*a, b, c*<br>*$\alpha, \beta, \gamma$* | 122.28,<br>122.28,<br>46.43,<br>90, 90,<br>120 | 122.28,<br>122.28,<br>46.43,<br>90, 90, 120 | 122.28,<br>122.28,<br>46.43,<br>90, 90, 120 | 122.3, 122.3,<br>46.45, 90, 90,<br>120 | 122.28,<br>122.28,<br>46.43,<br>90, 90, 120 | 122.28, 122.28,<br>46.43,<br>90, 90, 120 |
| Indexed patterns/indexing rate (%) | 68,252/<br>51.37 | 84,498/<br>40.63 | 102,276/<br>36.71 | 405789/<br>58.55 | 82,245/<br>41.30 | 105,277/<br>41.58 |
| Resolution range | 12.83-1.38<br>(1.48-1.38) | 12.75 - 1.51<br>(1.56 -<br>1.51) | 12.83 – 1.44<br>(1.50-1.44) | 19.42 – 1.85<br>(6.51-1.85) | 12.80 –<br>1.52 (1.57-1.52) | 12.88 - 1.49<br>(1.54-1.49) |
| No. of unique reflection | 55,868<br>(4000) | 51,980<br>(3387) | 54,980 (3820) | 35,025 (1828) | 51,170<br>(3302) | 52,115 (3524) |
| Completeness (%) | 100.00<br>(100.00) | 100.00<br>(100.00) | 100.00<br>(100.00) | 100.00 (100.00) | 100.00<br>(100.00) | 100.00 (100.00) |
| Redundancy | 3504<br>(652) | 975 (386) | 1340 (381) | 3266 (2739) | 1269 (394) | 1554 (421) |
| $R_{\text{split}}$ (%) | 3.9 (164) | 6.5<br>(144) | 6.4 (162) | 0.281 (1.20) | 7.1 (139) | 5.5 (149) |
| $I/\sigma(I)$ | 15.22<br>(0.64) | 9.76 (0.76) | 10.57 (0.66) | 8.32 (0.14) | 9.61 (0.76) | 11.28 (0.74) |
| $\text{CC}_{1/2}$ (%) | 0.997<br>(0.317) | 0.993<br>(0.300) | 0.993 (0.304) | 0.921 (0.615) | 0.992<br>(0.295) | 0.994 (0.292) |
| **Refinement statistics** |  |  |  |  |  |  |
| Estimated active state contribution | N/A | 8% | 10% | 10% | 8% | 7% |
| Refinement resolution ($\text{\AA}$) | 11.15-1.38<br>(1.41-1.38) | 10.93 –<br>1.80 (1.91 -<br>1.80) | 10.98 – 1.95<br>(2.10 – 1.95) | 19.40 - 2.30<br>(2.44 – 2.30) | 5.0 – 1.70<br>(1.79-170) | 10.88 – 1.70<br>(1.78-1.70) |
| No. reflections | 40,195<br>(2631) | 17,047<br>(2605) | 13,573 (2401) | 7,763 (1177) | 19,356<br>(2564) | 20,501 (2413) |
| $R_{\text{Work}} / R_{\text{Free}}$ | 0.1840 /<br>0.2014 | 0.3278 /<br>0.3368 | 0.2993 /<br>0.3402 | 0.3025 / 0.3297 | 0.3099 /<br>0.3480 | 0.3088 / 0.3569 |
| No. atoms | 1021 | 1074 | 1016 | 938 | 1012 | 1109 |
| Protein | 867 | 922 | 910 | 844 | 903 | 924 |
| Ligand | 31 | 31 | 31 | 31 | 31 | 31 |
| Solvent | 123 | 121 | 75 | 63 | 78 | 154 |
| Average atomic $B$-factor ($\text{\AA}^2$) | 36.86 | 24.46 | 42.19 | 25.7 | 15.07 | 11.67 |
| Protein | 35.10 | 22.93 | 41.49 | 25.73 | 14.22 | 10.15 |
| Ligand | 28.14 | 18.67 | 38.17 | 21.28 | 9.45 | 6.19 |
| Solvent | 51.51 | 37.64 | 52.24 | 27.49 | 27.14 | 21.88 |
| R.m.s. deviations |  |  |  |  |  |  |
| Bond lengths (Å) | 0.012 | 0.008 | 0.009 | 0.002 | 0.008 | 0.007 |
| Bond angles (°) | 1.27 | 1.10 | 1.20 | 0.54 | 1.01 | 1.01 |
| PDB code | 30NO | 30ML | 30MI | 30ZY | 30MK | 30QF |
| *Values in parentheses are for highest-resolution shell. |  |  |  |  |  |  |

| Data collection parameters | 100ns | 500ns | 1µs | 10µs | 20µs | 100µs |
| --- | --- | --- | --- | --- | --- | --- |
| Photon energy | 12.1 keV | 12.1 keV | 12.1 keV | 12.1 keV | 12.1 keV | 12.1 keV |
| Temperature (K) | 293 | 293 | 293 | 293 | 293 | 293 |
| X-ray beam size | 3x5 µm | 3x5 µm | 3x5 µm | 3x5 µm | 3x5 µm | 3x5 µm |
| Pulse energy | 580 µJ | 580 µJ | 580 µJ | 580 µJ | 580 µJ | 580 µJ |
| Pulse length | 25-30 fs | 25-30 fs | 25-30 fs | 25-30 fs | 25-30 fs | 25-30 fs |
| Repetition rate (X-ray/Laser) | 100/100 Hz | 100/100 Hz | 100/100 Hz | 100/100 Hz | 100/100 Hz | 100/100 Hz |
| Laser wavelength | 475 nm | 475 nm | 475 nm | 475 nm | 475 nm | 475 nm |
| Nozzle size | 75 µm | 75 µm | 75 µm | 75 µm | 75 µm | 75 µm |
| Jet speed | 28.1 mm s<sup>-1</sup> | 28.1 mm s<sup>-1</sup> | 28.1 mm s<sup>-1</sup> | 28.1 mm s<sup>-1</sup> | 28.1 mm s<sup>-1</sup> | 28.1 mm s<sup>-1</sup> |
| Data reduction |  |  |  |  |  |  |
| Space group | P 65 2 2 |  |  |  |  |  |
| Unit cell<br>*a, b, c*<br>*α, β, γ* | 122.28, 122.28, 46.43, 90, 90, 120 | 122.28, 122.28, 46.43, 90, 90, 120 | 122.28, 122.28, 46.43, 90, 90, 120 | 122.28, 122.28, 46.43, 90, 90, 120 | 122.28, 122.28, 46.43, 90, 90, 120 | 122.28, 122.28, 46.43, 90, 90, 120 |
| Indexed patterns/indexing rate (%) | 102,362/38.69 | 98,965/31.49 | 88,481/40.44 | 91,952/38.58 | 87,167/45.44 | 75,079/40.53 |
| Resolution range | 12.84 – 1.44 (1.50 -1.44) | 12.83 - 1.50 (1.55-1.50) | 12.85 – 1.51 (1.56-1.51) | 12.75-1.51 (1.56-1.51) | 12.84-1.52 (1.57-1.52) | 12.71-1.48 (1.53-1.48) |
| No. of unique reflection | 55,197 (3820) | 51,350 (3448) | 51,627 (3387) | 50,896 (3387) | 50,591 (3302) | 52,829 (3592) |
| Completeness (%) | 100.00 (100.00) | 100.00 (100.00) | 100.00 (100.00) | 100.00 (100.00) | 100.00 (100.00) | 100.00 (100.00) |
| Redundancy | 1641 (443) | 1391 (428) | 1157 (431) | 1247 (443) | 1186(466) | 1180 (418) |
| *R*<sub>split</sub> (%) | 6.1 (173) | 5.8 (129) | 6.1 (139) | 6.1 (131) | 6.4 (127) | 6.8 (153) |
| *I*/*σ*(*I*) | 10.46 (0.62) | 11.30 (0.87) | 10.36 (0.80) | 10.65 (0.83) | 10.16 (0.85) | 9.63 (0.70) |
| *CC*<sub>1/2</sub> | 0.994 (0.294) | 0.995 (0.326) | 0.995 (0.314) | 0.994 (0.344) | 0.993 (0.348) | 0.993 (0.311) |
| Wilson *B*-factor | 22.62 | 7.32 | 7.14 | 7.90 | 8.98 | 19.65 |
| Refinement statistics |  |  |  |  |  |  |
| Estimated active state contribution | 5% | 6% | 5% | 7% | 6% | 7% |
| Refinement resolution (Å) | 5.00 – 1.90 (2.04 -1.90) | 10.93 – 1.70 (1.79-1.70) | 5.00 – 1.80 (1.91-1.80) | 8.62 – 1.75 (1.84-1.75) | 7.0 – 1.75 (1.84-1.75) | 9.59-1.85 (1.96-1.85) |
| No. reflections | 11,704 (1896) | 19,568 (2675) | 15,923 (2533) | 18,574 (2516) | 17,842 (2379) | 15,193 (2345) |
| *R*<sub>Work</sub> / *R*<sub>Free</sub> | 0.4081 / 0.4400 | 0.3465 / 0.3851 | 0.3492 / 0.3755 | 0.3181 / 0.3451 | 0.3489 / 0.3567 | 0.3563 / 0.3944 |
| No. atoms | 1027 | 1074 | 1117 | 1008 | 1029 | 981 |
| Protein | 876 | 914 | 942 | 881 | 890 | 875 |
| Ligand | 31 | 31 | 62 | 31 | 31 | 31 |
| Solvent | 120 | 129 | 113 | 96 | 108 | 75 |
| Average atomic *B*-factor (Å<sup>2</sup>) | 28.42 | 14.60 | 11.45 | 14.80 | 16.71 | 26.44 |
| Protein | 27.86 | 12.95 | 10.68 | 13.65 | 15.46 | 25.73 |
| Ligand | 25.76 | 11.07 | 7.56 | 9.59 | 10.38 | 24.37 |
| Solvent | 33.07 | 27.15 | 19.99 | 27.07 | 28.80 | 35.68 |
| R.m.s deviations |  |  |  |  |  |  |
| Bond lengths (Å) | 0.005 | 0.007 | 0.011 | 0.009 | 0.009 | 0.01 |
| Bond angles (°) | 0.81 | 1.29 | 1.28 | 1.06 | 1.10 | 1.09 |
| PDB code | 30MM | 30MJ | 30ST | 30SE | 30SU | 30SV |
| *Values in parentheses are for highest-resolution shell. |  |  |  |  |  |  |

## Supplementary Text

The C4=O stretching mode serves as a reporter on the electronic state of the FMN during the photocycle. The followed frequencies correspond to the flavin dark state and the Cys-adduct (Extended Data Fig. 7a). Because of the width of infrared bands, contributions from the tail of nearby bands (e.g., amide I modes or other cofactor bands) may also be present at the selected frequencies. The corresponding time traces (Extended Data Fig. 7b) were fitted with a multiexponential function to model following events: dark state recovery from back reaction after flavin photoexcitation (τ_1_)^66^, Cys-adduct formation (τ_2_,τ_3_)^14^, changes in secondary structure after covalent bond formation (τ_4_, τ_5_)^67^.

Hydration levels of samples for time-resolved infrared spectroscopy were confirmed via the IR absorbance spectrum recorded with FT-IR spectroscopy (Extended Data Fig. 6). For this control experiment less protein volume was used, because this method requires less optical thickness of the sample than when measuring with the quantum cascade laser. Prominent amide I, amide II, and water bands prove that this procedure secures a well-hydrated sample.

**Movie S1. Structural dynamics of *Cr*LOV1 activation.** The movie shows the initial reaction of the flavin chromophore to the absorption of the photon, followed by the hydration of the chromophore environment essential for the key proton transfer and capture of the characteristic thioether bond formation in the LOV domain. It finally shows the rearrangement of the hydrogen bond network of the FMN environment involved in downstream signal transduction. The movie was generated using ChimeraX 1.10^68^ by a simple linear interpolation of coordinates between the 10 refined TR-SFX structures from 10 ps to 100 µs. The water entering the protein core is depicted as a red sphere.

---

**Supplementary Fig. 1:**
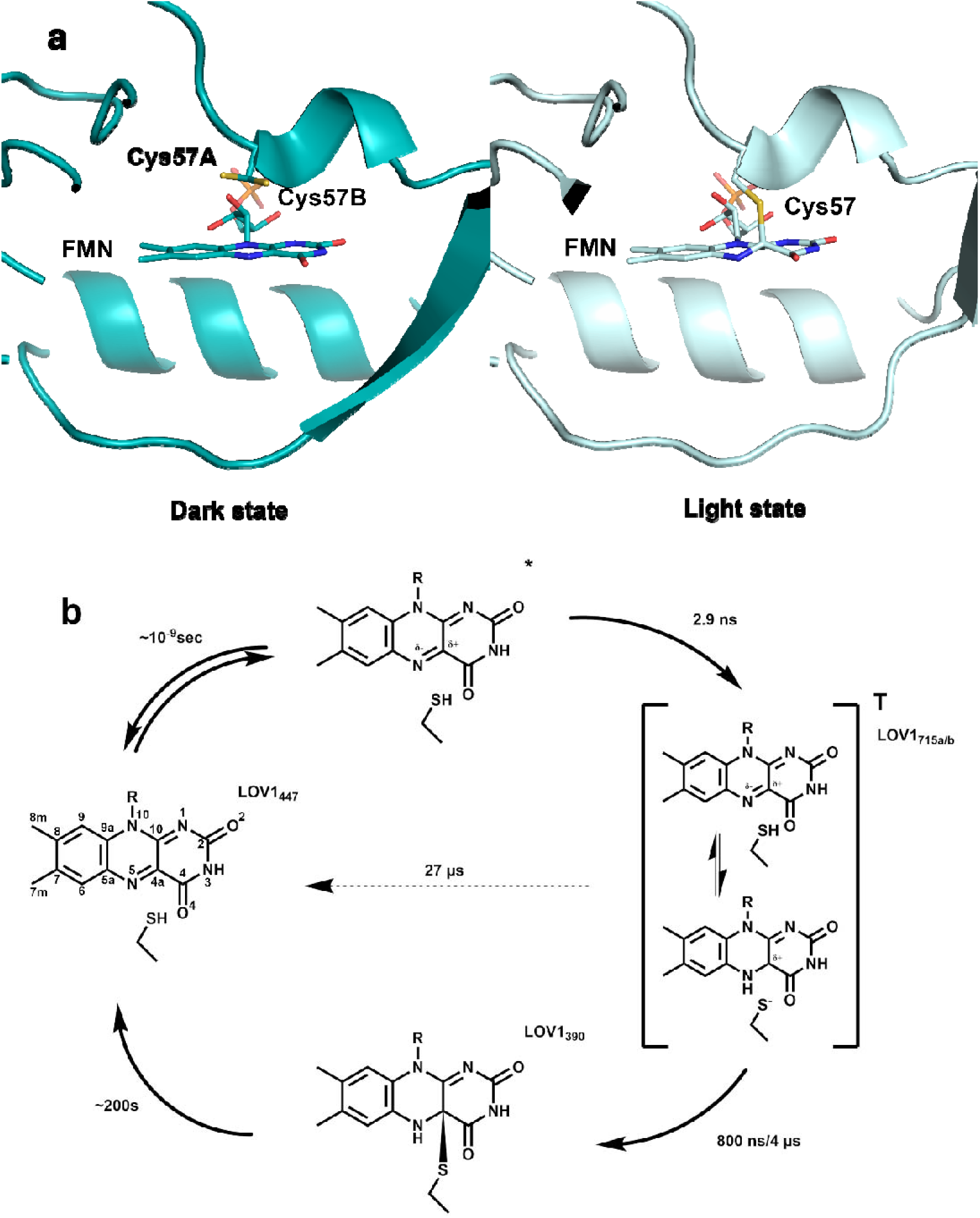
The thiol adduct formation of *Cr*PhotLOV1 upon light activation (A) and the photocycle as determined by Holzer et al. (2002)^13^ and Kottke et al. (2003)^14^ (B). **A)** The dark state structure (PDB: 8QI8) is shown as teal, the light state structure (PDB: 8QIA) as light blue cartoon^16^. The FMN and reactive cysteine (in alternate conformations) are shown in a stick representation. Residues 98-104 and 114-121 are hidden for better visibility. **B)** LOV1_447_ dark resting state forms a singlet excited state species upon light excitation (λ=447 nm). The singlet excited state decays into a triplet state intermediate (LOV1_715a/b_) via intersystem crossing. The protonation of the N5 position via the transfer of a hydrogen from the thiol group is thought to be possible. The reactive triplet state decays into the thiol adduct form, generating a long-lived light state species ^13,14^.

**Supplementary Fig. 2:**
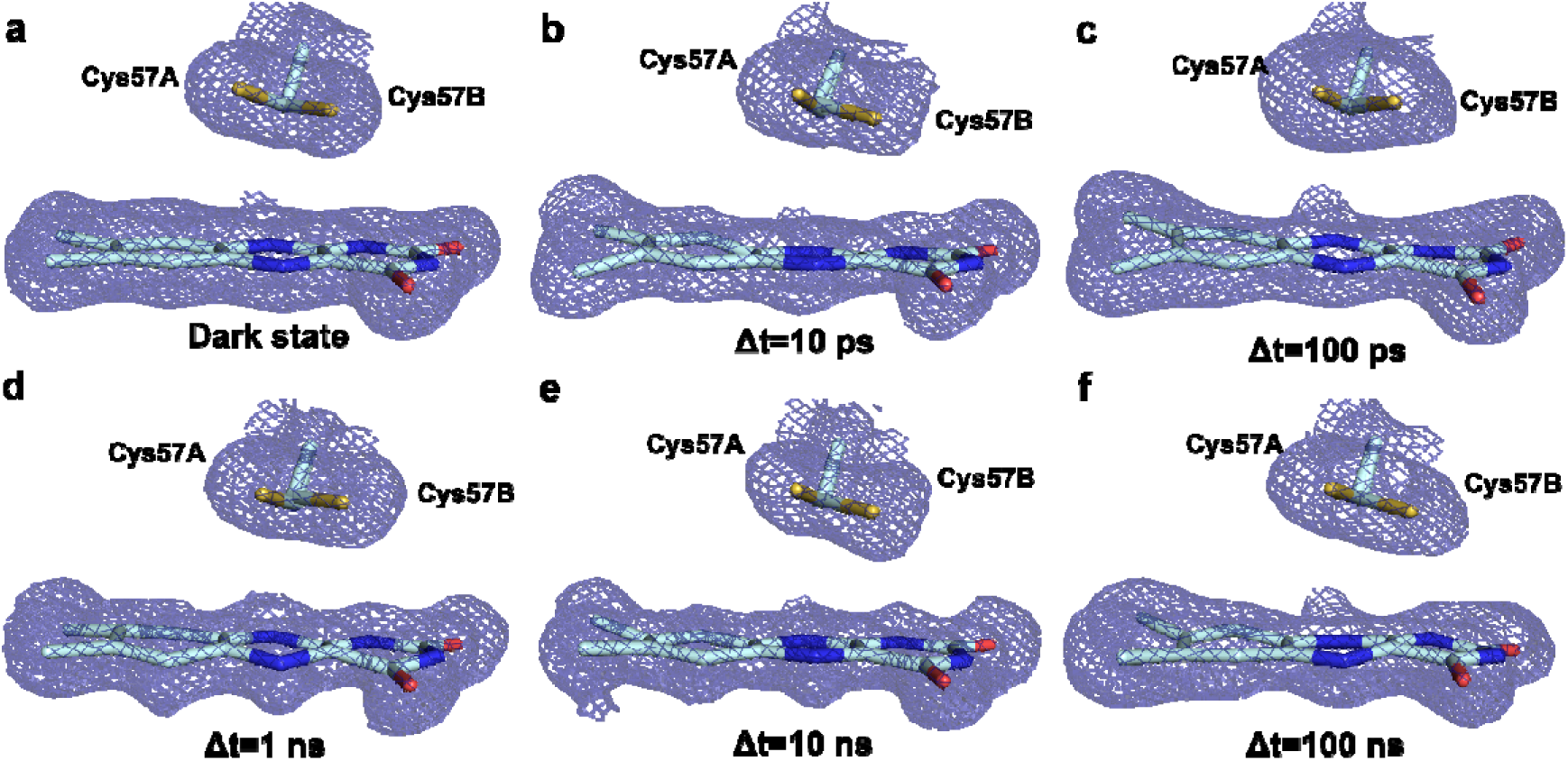
Distortions in the FMN ring structure with corresponding 2F_extrap_-F_c_ electron density. **A)** Dark resting state, **B)-D)** Δt=10 ps/100 ps/1 ns timepoints, corresponding to excited singlet state intermediates. **E)-F)** 10 ns/100 ns timepoints, corresponding to triplet state intermediates. The chromophore of the resting state and reaction intermediates are shown in light blue sticks, the reactive cysteine sidechain shown in identical representation, with alternate conformations labelled as A/B. The 2F_extrapol_-F_C_ electron density for the dark state, and the extrapolated electron density for the intermediates are shown as dark blue mesh, contoured at +1.0 σ around the shown residues.

**Supplementary Fig. 3:**
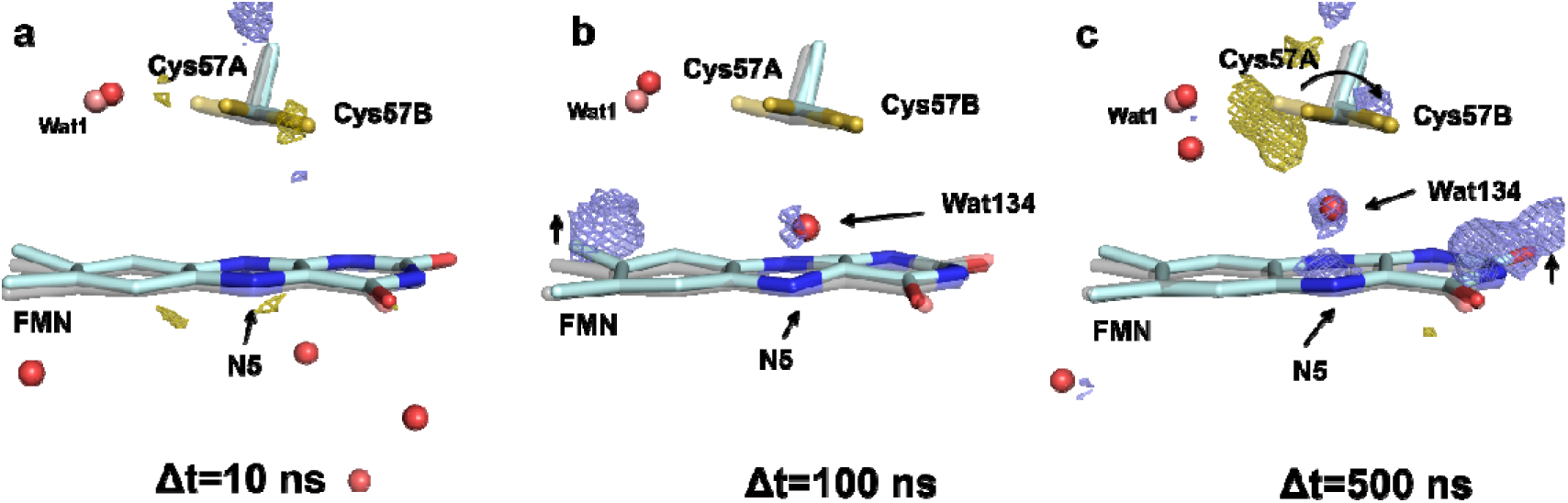
Hydration of the chromophore environment at the nanosecond timescale. **A)** At Δt=10 ns timepoint, corresponding to the transition to the triplet state, the FMN and reactive cysteine residue show similar position to the dark resting state. **B)** A transient water molecule appears at 100 ns post photoexcitation between the reactive cysteine and the N5 atom of FMN. **C)** The transient water occupancy increases at Δt=500 ns, as indicated by a positive difference density peak. The reactive cysteine begins to undergo a population redistribution, shifting towards the reactive B conformation. The structures of the reaction intermediates are shown in light blue sticks, overlaid with the dark resting state structure in grey transparent sticks. The reactive cysteine sidechains are shown in identical representation. The transient water Wat134 is shown as a red sphere (dark state water as salmon coloured spheres). The F_obs_^light^ _–_ F_obs_^dark^ difference Fourier electron density map contoured at ±3.0 σ, with positive density shown as blue, and negative as gold coloured mesh. Alternate conformations of the reactive Cys57 are labelled as A/B.

**Supplementary Fig. 4:**
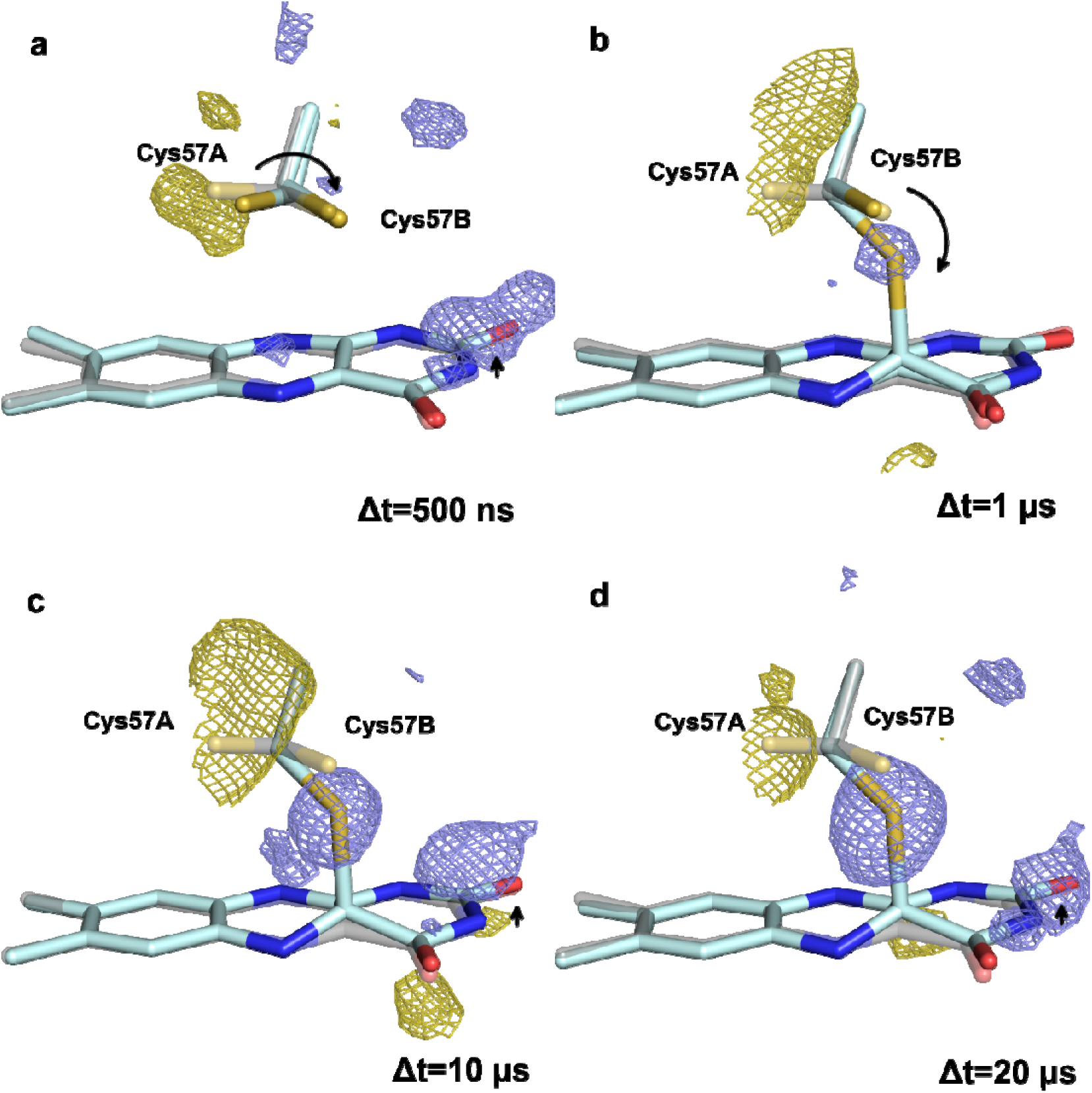
Formation of the covalent bond at the late nanosecond and early µs timescale is supported by the F_obs_^light^ _–_ F_obs_^dark^ difference density maps. **A**) In the final triplet state intermediate (Δt=500 ns) a shift in the Cys57 population distribution is present as indicated by the difference density peaks. The formation of the covalent bond is accompanied by the elevation of the pyrimidine ring, which begins at this stage. **B)** At the Δt=1 µs timepoint, the cysteine conformation A completely disappears, and the covalent bond formation begins. **C)-D)** At the subsequent intermediates (10 µs/20 µs post photoexcitation) the covalent bond is fully formed, and both Cys57 resting state conformations have disappeared. The elevation of the pyrimidine side of the isoalloxazine ring is also observed. The structure of the reaction intermediates are shown in pale cyan sticks, overlaid with the dark resting state structure in grey transparent sticks. The reactive cysteine sidechains are shown in identical representation, with alternate conformations labelled as A/B. The F_obs_^light^ – F_obs_^dark^ difference Fourier electron density map contoured at ±3.0 σ, with positive density shown as blue, and negative as gold coloured mesh.

**Supplementary Fig. 5:**
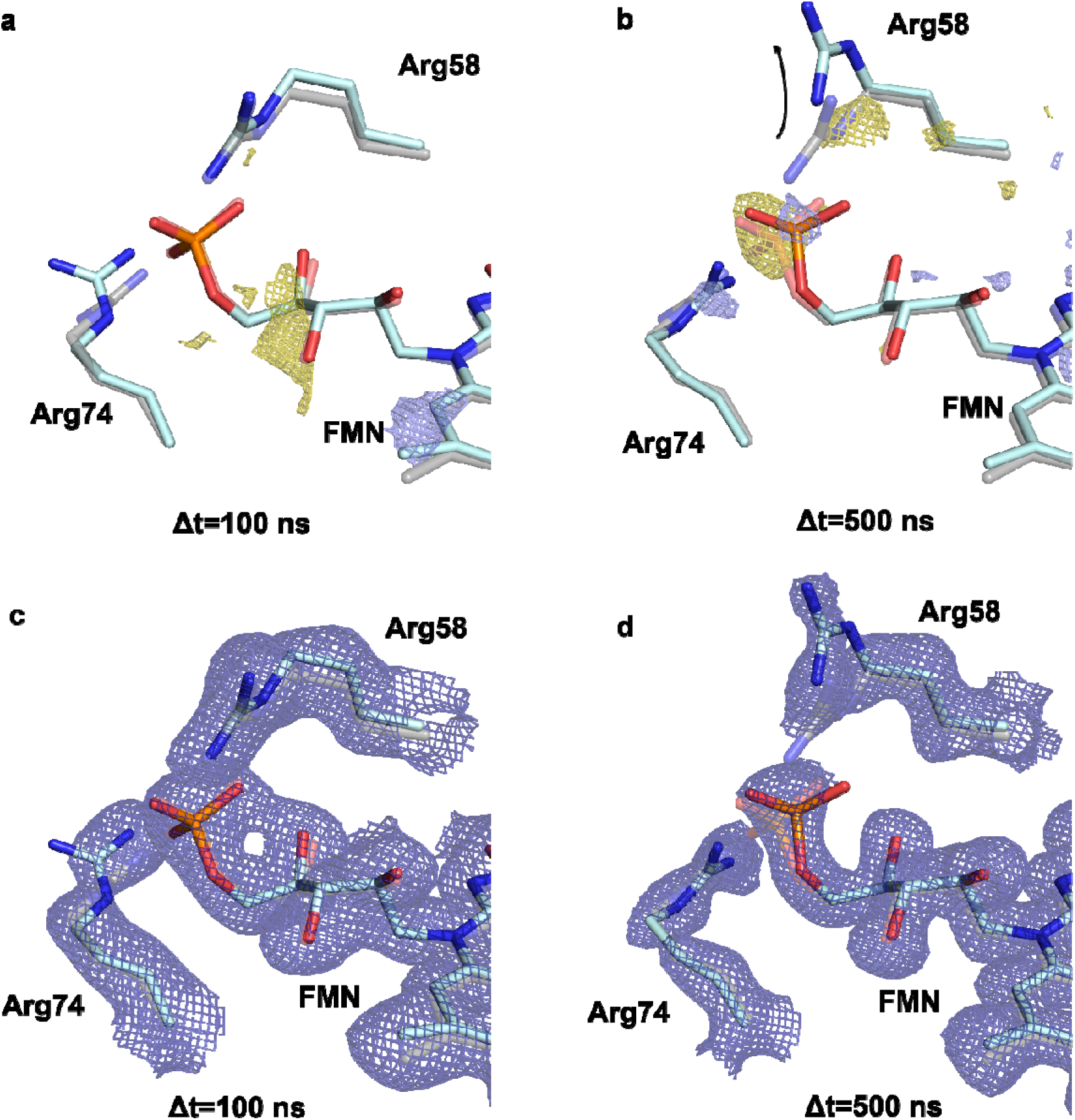
Displacement of the coordinating arginine residues accompanying the phosphate group disorder. **A)** The position of the FMN phosphate group, and the coordinating arginine residues are close to the resting state positions at the Δt=100 ns timepoint (triplet state intermediate) **B)** Along the shift of the phosphate group, both coordinating arginine residues shift in the Δt=500 ns intermediate, with Arg58 shifting away from the phosphate group, indicated by a strong negative difference density; and Arg74 moving closer to the resting state phosphate group position at this timepoint. **C)-D)** Panels showing the extrapolated electron density for the Δt=100 ns and Δt=500 ns intermediates. The F_light_ – F_dark_ difference Fourier electron density map contoured at ±3.0 σ, with positive density represented by blue, and negative density by gold coloured mesh. The 2F_extrapol_-F_C_ extrapolated electron density is shown as dark blue mesh, contoured at +1.0 σ around the residues depicted. The excited state chromophore is coloured light blue, overlaid with gray transparent sticks of the resting state structure.

**Supplementary Fig. 6:**
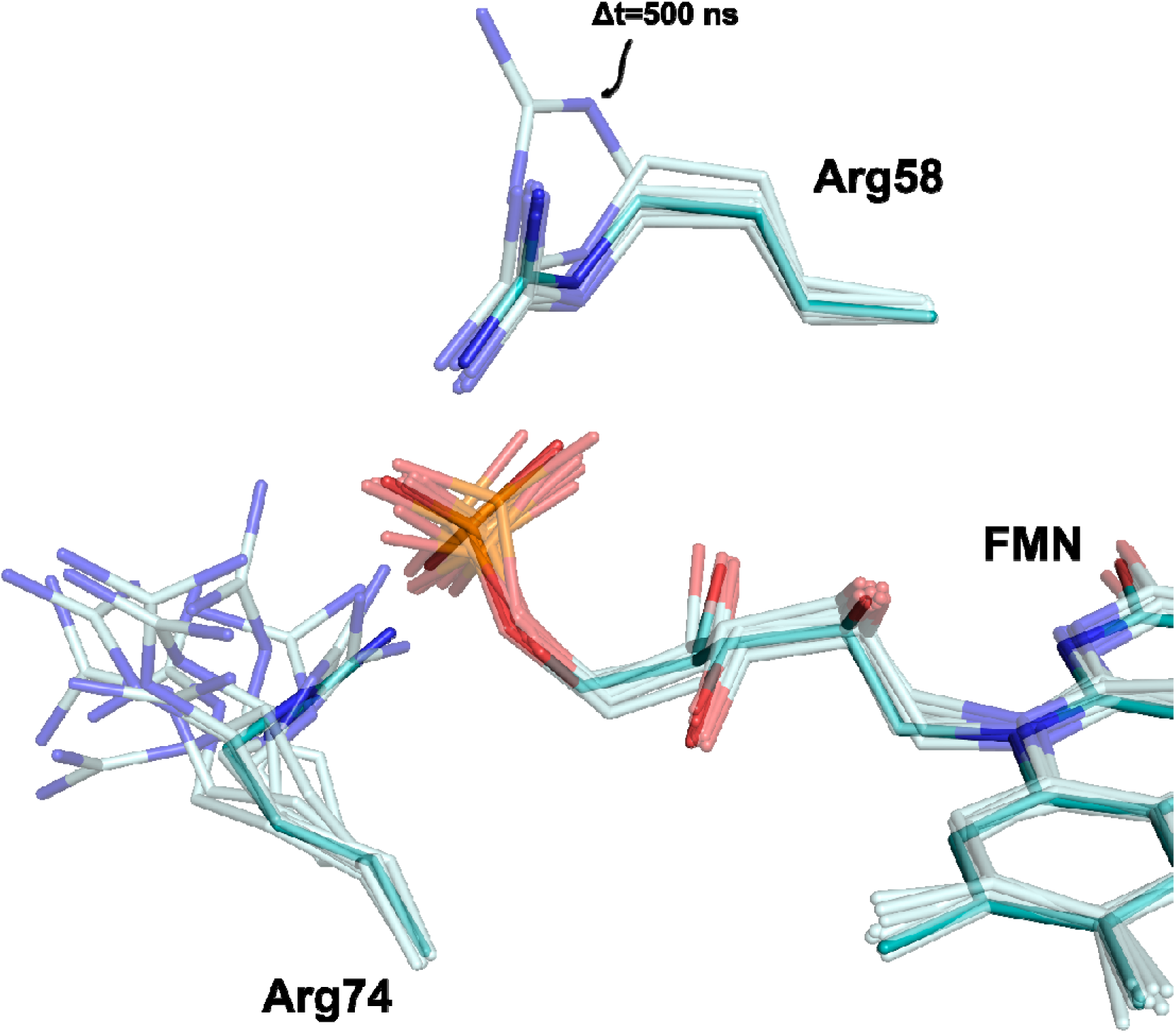
Increased mobility in the coordinating arginine residues in the intermediate structures. Teal sticks: The dark resting state positions of the coordinating arginine residues, Arg58 and Arg74. **Transparent, cyan sticks:** The increased mobility of the arginines observed in the probed intermediate structures, particularly prominent in the Arg74 position in the intermediate structures. Increased mobility in the Arg58 residue is accompanied by a disorder in the FMN phosphate group at the 500 ns timepoint.

**Supplementary Fig. 7:**
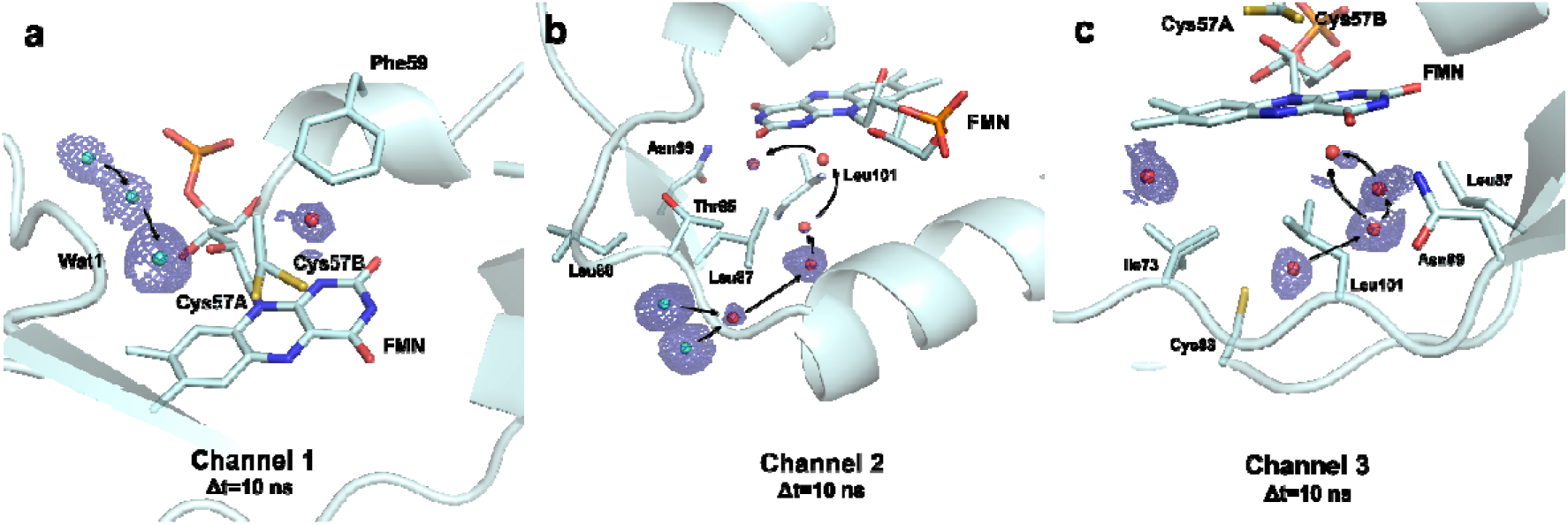
Hydration of protein core identified at the 10 ns intermediate. **A)** A set of water molecules present along the chromophore tail stricture leading to the resting state water closest to the reactive Cys57 residue, and a transient water near Phe59 appearing at the Δ*t*=10ps, 1 ns and 10 ns intermediate. **B)** A second water channel running from the surface to the chromophore binding pocket along Leu87, Thr65 and Leu101. **C)** A third transient water channel identified from the surface near Cys83 leading to the isoalloxazine ring along Asn99. The protein molecule is shown in light blue cartoon and sticks, water molecules are represented by red spheres. Waters present in the resting state are coloured cyan. The electron density is calculated from the extrapolated structure factors used for structure refinement, contoured at 0.8 σ 1.3 Å around the transient water molecules, and is shown as dark blue mesh. Alternate conformations of the reactive Cys57 are labelled as A/B.

**Supplementary Fig. 8:**
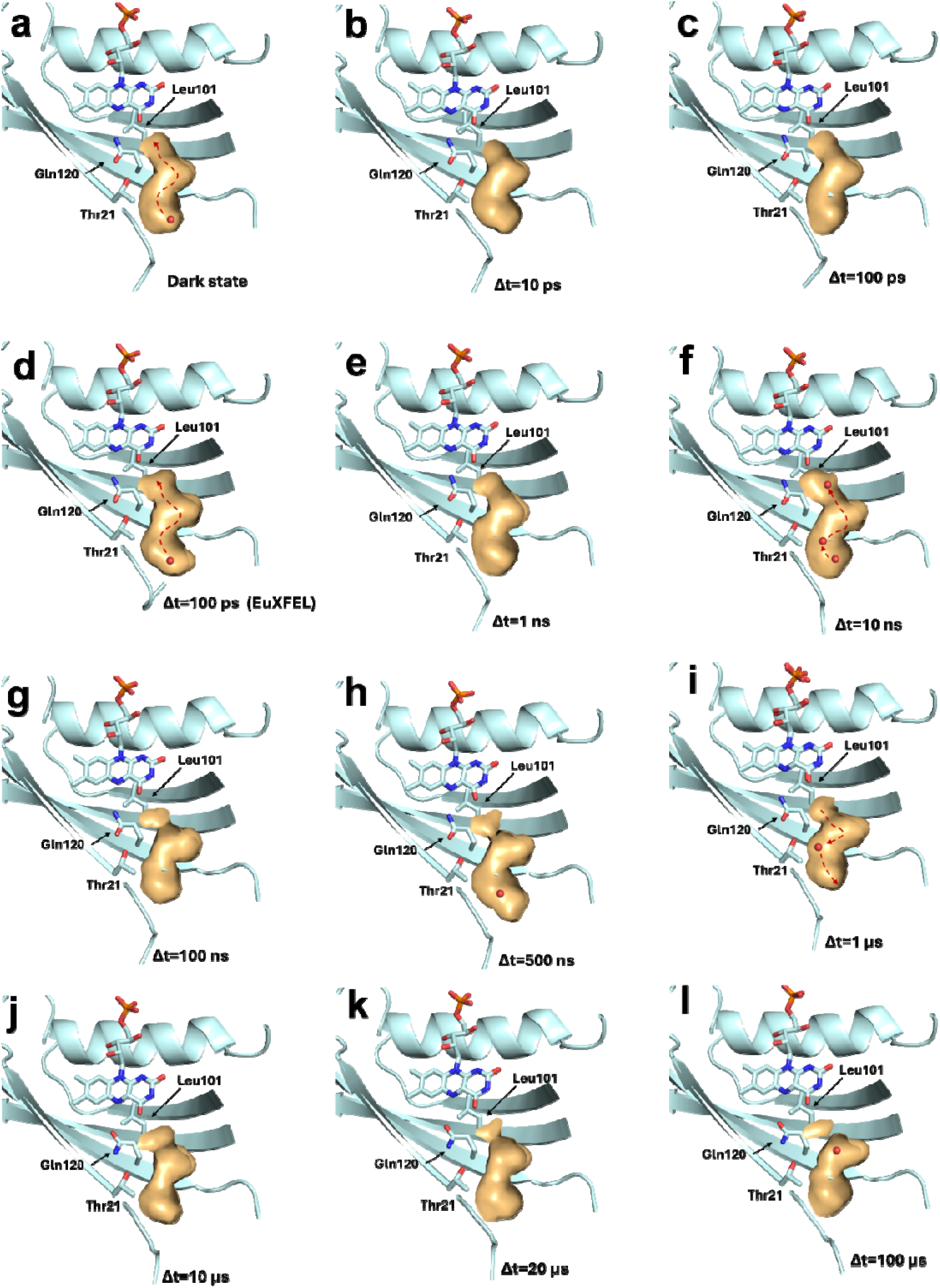
Time-resolved views of the CrLOV1 water channel across the photocycle. Structural snapshots at selected time points illustrate the position of the tunnel and associated water molecules, highlighting dynamic changes in solvent accessibility during light activation. Protein shown as teal cartoon, tunnel surface in orange, and water molecules as red spheres.

**Supplementary Fig. 9:**
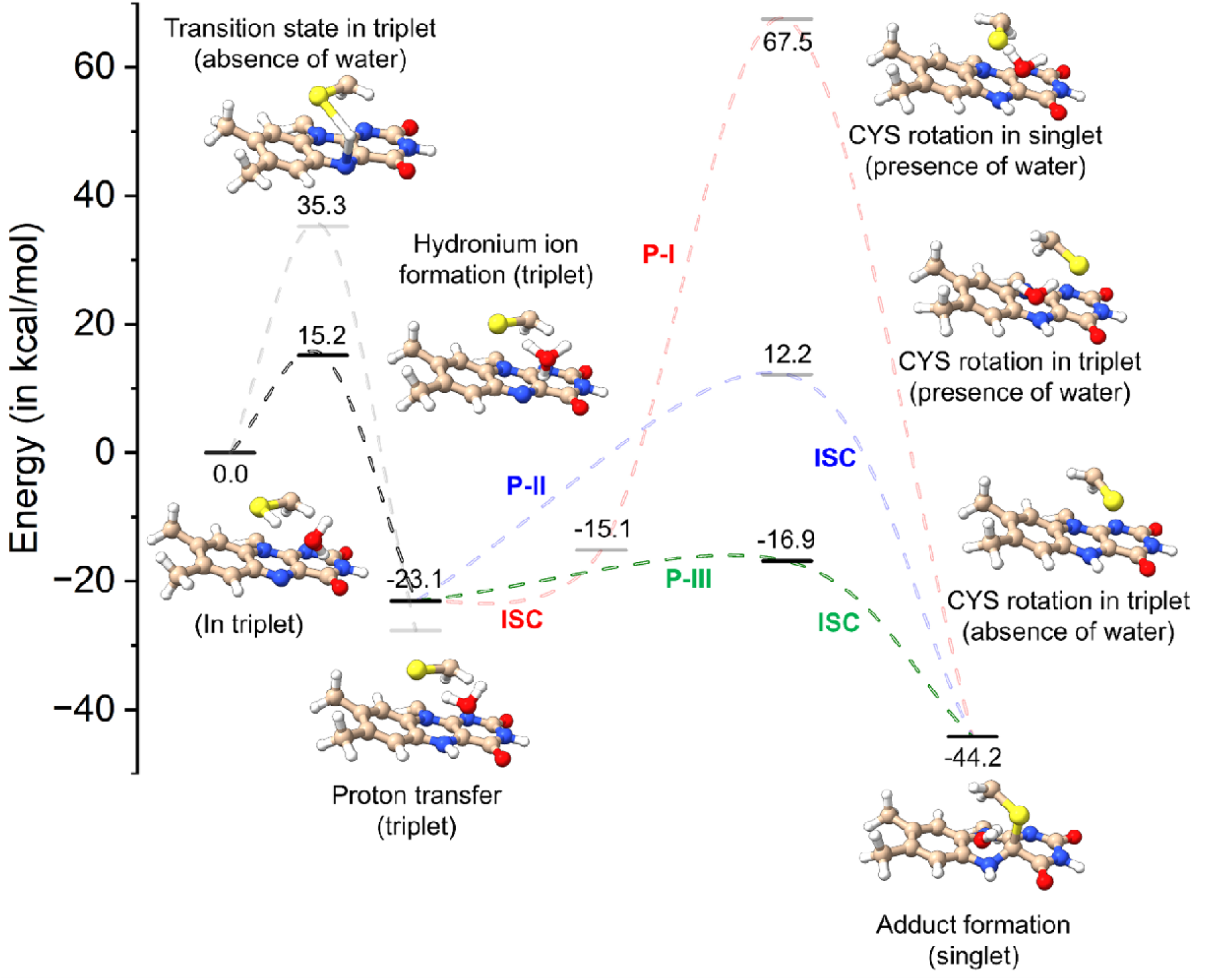
Energy profile of proton transfer and adduct formation taking place between Cys and FMN. The complete calculated QM/MM energy profile for the proton transfer and adduct formation pathway of the *Cr*LOV1 photocycle. The transition state in triplet is energetically favoured in the presence of water (Hydronium ion formation). The ISC pathway P-I depicts the Cys57 rotation in singlet state in the presence of water, P-II in triplet in the presence of water, and P-III in triplet state in the absence of water (most favourable pathway).

**Supplementary Fig. 10:**
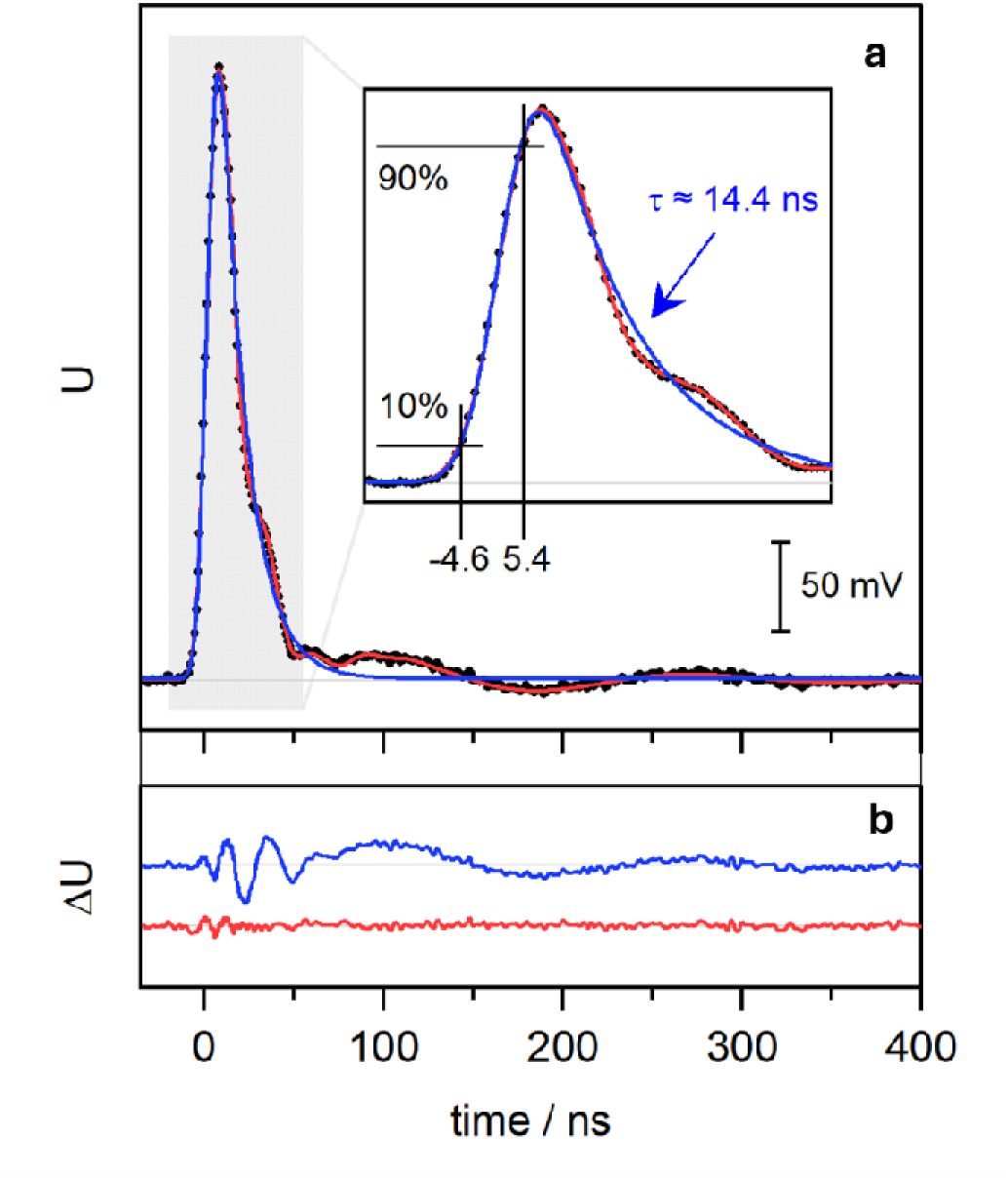
Instrument response measurement of the EC-QCL based time-resolved infrared spectroscopy setup. **A)** Single acquisition of the transient voltage changes after flashing the detection element of the sample MCT detector directly with a strongly attenuated 532 nm laser pulse (5 ns pulse width). The trace was fitted with an exponentially modified gaussian (EMG) function (blue) and the sum of an EMG and two dampened sine functions (red). **B)** Residuals of the fits shown in A using the same vertical scale and same colour code.

**Supplementary Fig. 11:**
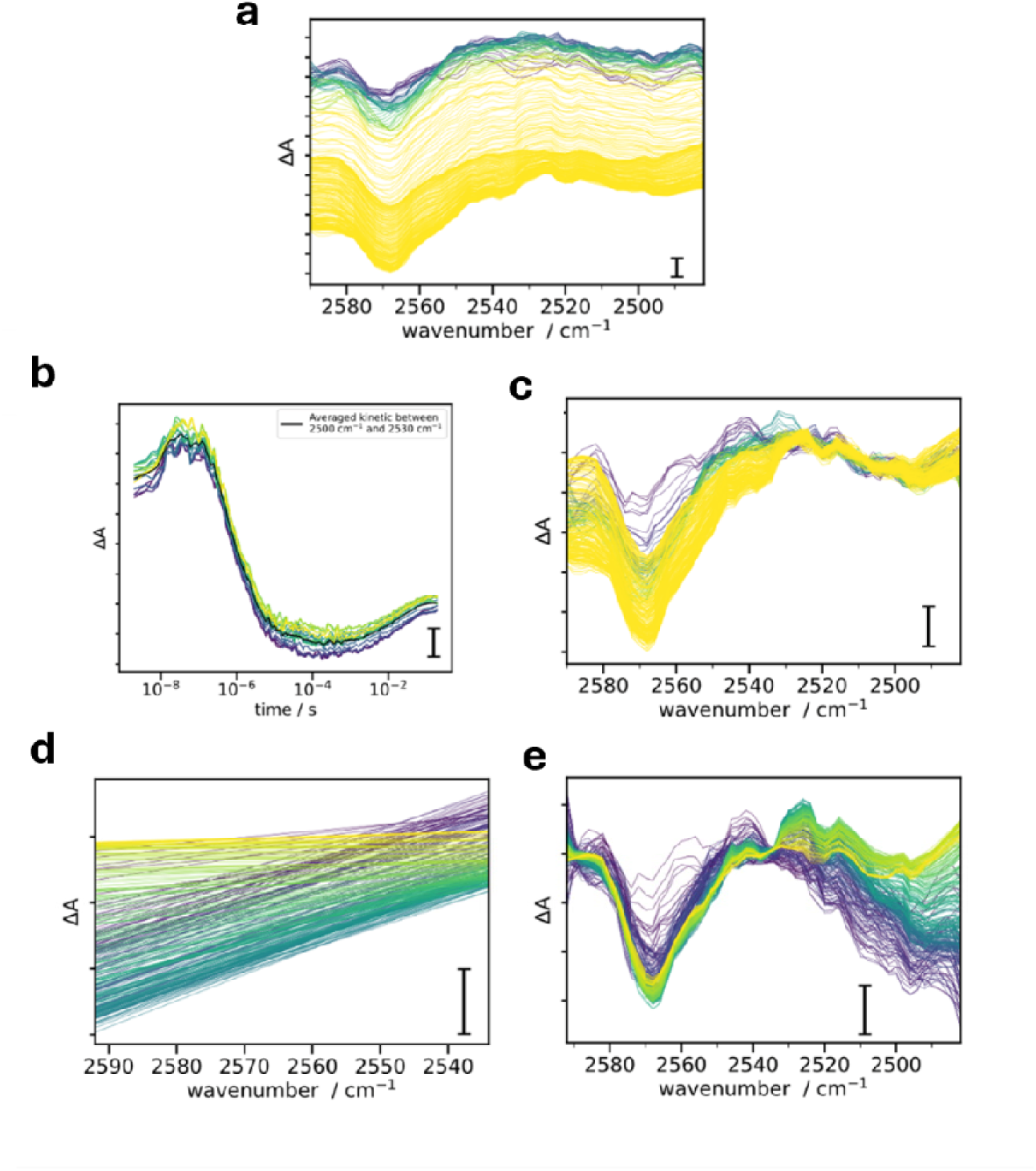
Heat signal removal of the time-resolved infrared spectroscopy dataset of the S-H stretching frequency range. **A)** Time-resolved IR difference spectra after smoothing with Savitsky-Golay filter. **B)** Averaged kinetic trace between 2530 and 2500 cm^-1^ to approximate major heating signal. **C)** Time-resolved IR difference spectra after subtracting averaged major heating signal. **D)** 1d polynomial fit between 2592 and 2533 cm^-1^ for each spectrum to approximate remaining minor heating signal. **E)** Time-resolved IR difference spectra after subtracting minor heating signal. For all the figures, the spectra were coloured purple to yellow from early times to late time. Scale bars, 500 µOD (See Materials & Methods for details on the analysis.)

## References

1 Kutta, R. J., Magerl, K., Kensy, U. & Dick, B. A search for radical intermediates in the photocycle of LOV domains. Photochem Photobiol Sci 14, 288–299, doi:10.1039/c4pp00155a (2015).

2 Briggs, W. R. et al. The phototropin family of photoreceptors. The Plant Cell 13, 993–997 (2001).

3 Li, F. W. et al. The origin and evolution of phototropins. Front Plant Sci 6, 637, doi:10.3389/fpls.2015.00637 (2015).

4 Sakai, T. et al. Arabidopsis nph1 and npl1: blue light receptors that mediate both phototropism and chloroplast relocation. PNAS 98, 6969–6974 (2001).

5 Kinoshita, T. et al. phot1 and phot2 mediate blue light regulation of stomatal opening. Nature 414, 656–660 (2001).

6 Kagawa, T. et al. Arabidopsis NPL1: A phototropin homolog controlling the chloroplast high light avoidance response. Science 291, 2138–2141 (2001).

7 Huang, K. & Beck, C. F. Phototropin is a blue-light receptor that controls multiple steps in the sexual life cycle of the green alga Chlamydomonas reinhardtii. PNAS 100, 6269–6274 (2003).

8 Im, C. S., Eberhard, S., Huang, K., Beck, C. F. & Grossman, A. R. Phototropin involvement in the expression of genes encoding chlorophyll and carotenoid biosynthesis enzymes and LHC apoproteins in Chlamydomonas reinhardtii. Plant J 48, 1–16, doi:10.1111/j.1365-313X.2006.02852.x (2006).

9 Petroutsos, D. et al. A blue-light photoreceptor mediates the feedback regulation of photosynthesis. Nature 537, 563–566, doi:10.1038/nature19358 (2016).

10 Trippens, J. et al. Phototropin influence on eyespot development and regulation of phototactic behavior in Chlamydomonas reinhardtii. Plant Cell 24, 4687–4702, doi:10.1105/tpc.112.103523 (2012).

11 Salomon, M. et al. An optomechanical transducer in the blue light receptor phototropin from Avena sativa. PNAS 98, 12357–12361 (2001).

12 Crosson, S. & Moffat, K. Structure of a flavin-binding plant photoreceptor domain: Insights into light-mediated signal transduction. PNAS 98, 2995–3000 (2001).

13 Holzer, W., Penzkofer, A., Fuhrmann, M. & Hegemann, P. Spectroscopic characterization of flavin mononucleotide bound to the LOV1 domain of Phot1 from Chlamydomonas reinhardtii. Photochem Photobiol 75, 479–487, doi: 10.1562/0031-8655(2002)075&<0479:scofmb>2.0.co;2 (2002).

14 Kottke, T., Heberle, J., Hehn, D., Dick, B. & Hegemann, P. Phot-LOV1: photocycle of a blue-light receptor domain from the green alga Chlamydomonas reinhardtii. Biophys J 84, 1192–1201 (2003).

15 Fedorov, R. et al. Crystal structures and molecular mechanism of a light-induced signaling switch: the phot-LOV1 domain from Chlamydomonas reinhardtii. Biophys J 84, 2474–2482 (2003).

16 Gotthard, G. et al. Capturing the blue-light activated state of the Phot-LOV1 domain from Chlamydomonas reinhardtii using time-resolved serial synchrotron crystallography. IUCrJ 11, 792–808, doi:10.1107/S2052252524005608 (2024).

17 Corchnoy, S. B. et al. Intramolecular proton transfers and structural changes during the photocycle of the LOV2 domain of phototropin 1. J Biol Chem 278, 724–731, doi:10.1074/jbc.M209119200 (2003).

18 Kennis, J. T. M. et al. Primary Reactions of the LOV2 Domain of Phototropin, a Plant Blue-Light Photoreceptor. Biochemistry 42, 3385–3392 (2003).

19 Lanzl, K., Noll, G. & Dick, B. LOV1 protein from Chlamydomonas reinhardtii is a template for the photoadduct formation of FMN and methylmercaptan. Chembiochem 9, 861–864, doi:10.1002/cbic.200700737 (2008).

20 Lanzl, K., Sanden-Flohe, M. V., Kutta, R. J. & Dick, B. Photoreaction of mutated LOV photoreceptor domains from Chlamydomonas reinhardtii with aliphatic mercaptans: implications for the mechanism of wild type LOV. Phys Chem Chem Phys 12, 6594–6604, doi:10.1039/b922408d (2010).

21 Zayner, J. P. & Sosnick, T. R. Factors that control the chemistry of the LOV domain photocycle. PLoS One 9, e87074, doi:10.1371/journal.pone.0087074 (2014).

22 Eyring, H. The activated complex in chemical reactions. Journal of Chemical Physics 3, 107–115 (1935).

23 Ataka, K., Hegemann, P. & Heberle, J. Vibrational spectroscopy of an algal Phot-LOV1 domain probes the molecular changes associated with blue-light reception. Biophys J 84, 466–474 (2003).

24 Iwata, T., Tokutomi, S. & Kandori, H. Photoreaction of the cysteine S-H group in the LOV2 domain of Adiantum phytochrome3. J Am Chem Soc 124, 11840–11841 (2002).

25 Iuliano, J. N. et al. Unraveling the Mechanism of a LOV Domain Optogenetic Sensor: A Glutamine Lever Induces Unfolding of the Jalpha Helix. ACS Chem Biol 15, 2752–2765, doi:10.1021/acschembio.0c00543 (2020).

26 Sato, Y., Iwata, T., Tokutomi, S. & Kandori, H. Reactive cysteine is protonated in the triplet excited state of the LOV2 domain in Adiantum Phyctochrome3. J Am Chem Soc 127, 1088–1089 (2005).

27. Maity, S. et al. Hydraulic Activation of the AsLOV2 Photoreceptor. bioRxiv, doi:10.1101/2025.06.19.660617 (2025).

## Methods references

28 Studier, F. W. Protein production by auto-induction in high density shaking cultures. Protein Expr Purif 41, 207–234, doi:10.1016/j.pep.2005.01.016 (2005).

29 Weierstall, U. et al. Lipidic cubic phase injector facilitates membrane protein serial femtosecond crystallography. Nat Commun 5, 3309, doi:10.1038/ncomms4309 (2014).

30 Weierstall, U. et al. Droplet streams for serial crystallography of proteins. Experiments in Fluids 44, 675–689, doi:10.1007/s00348-007-0426-8 (2007).

31 DePonte, D. P. et al. Gas dynamic virtual nozzle for generation of microscopic droplet streams. J. Phys. D: Appl. Phys. 41, 195505 (2008).

32 Vakili, M. et al. 3D printed devices and infrastructure for liquid sample delivery at the European XFEL. J Synchrotron Radiat 29, 331–346, doi:10.1107/S1600577521013370 (2022).

33 Mancuso, A. P. et al. The Single Particles, Clusters and Biomolecules and Serial Femtosecond Crystallography instrument of the European XFEL: initial installation. J Synchrotron Radiat 26, 660–676, doi:10.1107/S1600577519003308 (2019).

34 Koliyadu, J. C. P. et al. Pump-probe capabilities at the SPB/SFX instrument of the European XFEL. J Synchrotron Radiat 29, 1273–1283, doi:10.1107/S1600577522006701 (2022).

35 Wiedorn, M. O. et al. Megahertz serial crystallography. Nat Commun 9, 4025, doi:10.1038/s41467-018-06156-7 (2018).

36 Allahgholi, A. et al. AGIPD, a high dynamic range fast detector for the European XFEL. Journal of Instrumentation 10, C01023–C01023, doi:10.1088/1748-0221/10/01/c01023 (2015).

37 Hauf, S. et al. The Karabo distributed control system. J Synchrotron Radiat 26, 1448–1461, doi:10.1107/S1600577519006696 (2019).

38 Palmer, G. et al. Pump-probe laser system at the FXE and SPB/SFX instruments of the European X-ray Free-Electron Laser Facility. J Synchrotron Radiat 26, 328–332, doi:10.1107/S160057751900095X (2019).

39 Schultz, B. J., Mohrmann, H., Lorenz-Fonfria, V. A. & Heberle, J. Protein dynamics observed by tunable mid-IR quantum cascade lasers across the time range from 10ns to 1s. Spectrochim Acta A Mol Biomol Spectrosc 188, 666–674, doi:10.1016/j.saa.2017.01.010 (2018).

40 Maia, R. N. A. et al. Real-Time Tracking of Proton Transfer from the Reactive Cysteine to the Flavin Chromophore of a Photosensing Light Oxygen Voltage Protein. J Am Chem Soc 143, 12535–12542, doi:10.1021/jacs.1c03409 (2021).

41 White, T. A. et al. CrystFEL: a software suite for snapshot serial crystallography. Journal of Applied Crystallography 45, 335–341, doi:10.1107/s0021889812002312 (2012).

42 Gevorkov, Y. et al. XGANDALF - extended gradient descent algorithm for lattice finding. Acta Crystallogr A Found Adv 75, 694–704, doi:10.1107/S2053273319010593 (2019).

43 Powell, H. R. The Rossmann Fourier autoindexing algorithm in MOSFLM. Acta Cryst. D 55, 1690–1695 (1999).

44 White, T. A. et al. Recent developments in CrystFEL. J Appl Crystallogr 49, 680–689, doi:10.1107/S1600576716004751 (2016).

45 Turkot, O. et al. EXtra-Xwiz: A Tool to Streamline Serial Femtosecond Crystallography Workflows at European XFEL. Crystals 13, doi:10.3390/cryst13111533 (2023).

46 White, T. A. Processing serial crystallography data with CrystFEL: a step-by-step guide. Acta Crystallogr D Struct Biol 75, 219–233, doi:10.1107/S205979831801238X (2019).

47 Barty, A. et al. Cheetah: software for high-throughput reduction and analysis of serial femtosecond X-ray diffraction data. J Appl Crystallogr 47, 1118–1131, doi:10.1107/S1600576714007626 (2014).

48 Yefanov, O. et al. Accurate determination of segmented X-ray detector geometry. Opt Express 23, 28459–28470, doi:10.1364/OE.23.028459 (2015).

49 White, T. A. Detector alignment for X-ray crystallography using Millepede-II. J Appl Crystallogr 59, 594–608, doi:10.1107/S1600576726001287 (2026).

50 Mous, S. et al. Dynamics and mechanism of a light-driven chloride pump. Science 375, 845–851 (2022).

51 McCoy, A. J. et al. Phaser crystallographic software. J Appl Crystallogr 40, 658–674, doi:10.1107/S0021889807021206 (2007).

52 Emsley, P., Lohkamp, B., Scott, W. G. & Cowtan, K. Features and development of Coot. Acta Crystallogr D Biol Crystallogr 66, 486–501, doi:10.1107/S0907444910007493 (2010).

53 Liebschner, D. et al. Macromolecular structure determination using X-rays, neutrons and electrons: recent developments in Phenix. Acta Crystallogr D Struct Biol 75, 861–877, doi:10.1107/S2059798319011471 (2019).

54 Schneider, C. & Suhnel, J. A molecular dynamics simulation of the flavin mononucleotide-RNA aptamer complex. Biopolymers 50, 287–302, doi: 10.1002/(sici)1097-0282(199909)50:3&<287::Aid-bip5>3.0.Co;2-g (1999).

55 Case, D. A. et al. AmberTools. J Chem Inf Model 63, 6183–6191, doi:10.1021/acs.jcim.3c01153 (2023).

56. Jonsson, H., Mills, G. & Jacobsen, K. W. in Classical and quantum dynamics in condensed phase simulations (eds B. J. Berne, G. Ciccotti, & D. F. Coker) Ch. 16, 385–404 (1998).

57 Yanai, T., Tew, D. P. & Handy, N. C. A new hybrid exchange–correlation functional using the Coulomb-attenuating method (CAM-B3LYP). Chemical Physics Letters 393, 51–57, doi:10.1016/j.cplett.2004.06.011 (2004).

58 Grimme, S., Antony, J., Ehrlich, S. & Krieg, H. A consistent and accurate ab initio parametrization of density functional dispersion correction (DFT-D) for the 94 elements H-Pu. J Chem Phys 132, 154104, doi:10.1063/1.3382344 (2010).

59 Dunning Jr., T. H. Gaussian basis sets for use in correlated molecular calculations. I. The atoms boron through neon and hydrogen. J. Chem. Phys 90, 1007–1023 (1988).

60 Maier, J. A. et al. ff14SB: Improving the Accuracy of Protein Side Chain and Backbone Parameters from ff99SB. J Chem Theory Comput 11, 3696–3713, doi:10.1021/acs.jctc.5b00255 (2015).

61 Jorgensen, W. L., Chandrasekhar, J. & Madura, J. D. Comparison of simple potential functions for simulating liquid water. J. Chem. Phys 79, 926–935 (1983).

62 Ufimtsev, I. S. & Martinez, T. J. Quantum Chemistry on Graphical Processing Units. 3. Analytical Energy Gradients, Geometry Optimization, and First Principles Molecular Dynamics. J. Chem. Theory Comput. 5, 2619–2628 (2009).

63 Titov, A. V., Ufimtsev, I. S., Luehr, N. & Martinez, T. J. Generating Efficient Quantum Chemistry Codes for Novel Architectures. J Chem Theory Comput 9, 213–221, doi:10.1021/ct300321a (2013).

64 Kastner, J. et al. DL-FIND: an open-source geometry optimizer for atomistic simulations. J. Phys. Chem. 113, 11856–11865 (2009).

65 Metz, S., Kästner, J., Sokol, A. A., Keal, T. W. & Sherwood, P. ChemShell—a modular software package for QM/MM simulations. WIREs Computational Molecular Science 4, 101–110, doi:10.1002/wcms.1163 (2013).

## Supplementary Information References

66 Alexandre, M. T. A. et al. Primary reactions of the LOV2 domain of phototropin studied with ultrafast mid-infrared spectroscopy and quantum chemistry. Biophys J 97, 227–237 (2009).

67 Pfeifer, A. et al. Time-resolved Fourier transform infrared study on photoadduct formation and secondary structural changes within the phototropin LOV domain. Biophys J 96, 1462–1470, doi:10.1016/j.bpj.2008.11.016 (2009).

68 Pettersen, E. F. et al. UCSF ChimeraX: Structure visualization for researchers, educators, and developers. Protein Sci 30, 70–82, doi:10.1002/pro.3943 (2021).

